# Rapid Human Oogonia-like Cell Specification via Combinatorial Transcription Factor-Directed Differentiation

**DOI:** 10.1101/2022.07.11.499564

**Authors:** Merrick Pierson Smela, Christian C Kramme, Patrick R.J. Fortuna, Bennett Wolf, Venkata Srikar Kavirayuni, Jessica Adams, Carl Ma, Sergiy Velychko, Ursula Widocki, Shrey Goel, Tianlai Chen, Sophia Vincoff, Edward Dong, Richie E. Kohman, Mutsumi Kobayashi, Toshi Shioda, George M. Church, Pranam Chatterjee

## Abstract

The generation of germline cell types from human induced pluripotent stem cells (hiPSCs) represents a key milestone toward *in vitro* gametogenesis, which has the potential to transform reproductive modeling and medicine. Methods to recapitulate advanced germline cell specification *in vitro* have relied on extensive, long term culture methods, the most notable of which is a four-month culture protocol employing xenogeneic reconstituted ovaries with mouse embryonic ovarian somatic cells. Recently, transcription factor (TF)-based methods have demonstrated the feasibility of exogenous factor expression to directly differentiate hiPSCs into cell types of interest, including various ovarian cell types. The protocols leveraged in these studies, however, utilize more local methods of factor selection, such as basic differential gene expression analysis, and lower-throughput screening strategies via iterative testing of a small set of TFs. In this work, we integrate our recently-described graph theory pipeline and highly-parallelized screening protocols to globally identify and screen 46 oogenesis-regulating TFs for their role in human germline formation. We identify *ZNF281*, *LHX8*, *SOHLH1, ZGLP1,* and *ANHX* whose combinatorial overexpression drives DDX4+ induced oogonia-like cell (iOLC) formation from hiPSCs. In contrast to previous methods, our protocol employs a simple four-day, feeder-free monolayer culture condition. We additionally demonstrate a method of post-isolation, feeder-free expansion of DDX4+ iOLCs that shows retained cell identity *in vitro*. We additionally identify *DLX5*, *HHEX*, and *FIGLA* whose individual overexpression enhances hPGCLC formation from hiPSCs. We characterize these TF-based iOLCs and hPGCLCs via gene and protein expression analyses and demonstrate their broad similarity to *in vivo* and *in vitro*-derived oogonia and primordial germ cells. Together, these results identify new regulatory factors that enhance *in vitro* human germ cell specification and further establish unique computational and experimental tools for human *in vitro* oogenesis research.

## Main

In the human germline, human oogonia are specified from primordial germ cells (PGCs) that undergo sequential differentiation through sex-specific trajectories to form female gametes.^1^ *In vitro* gametogenesis (IVG) is a powerful technique for the recapitulation of germline development, often utilizing growth factor and co-culture directed differentiation of pluripotent stem cells to sequentially specify discrete germline cell types.^2^ Germ cells are challenging to obtain for research in humans due to inherent ethical and technical limitations in their *in vivo* isolation and utilization, since these critical intermediates span embryonic, fetal, and adult development stages.^3^ With the rate of infertility on the rise globally, new cell models of human reproductive development are needed to accelerate research into sex cell formation and genetic regulation, investigate root causes of infertility, and to develop novel assisted reproductive technologies.^4^

Methods for generating primordial germ cell-like cells (hPGCLCs), which highly resemble pre-migratory *in vivo* PGCs, from stem cells are abundant and utilize a variety of both three-dimensional and two-dimensional methodologies.^1,2,5–13^ Recently, human DDX4+ oogonia were specified from pluripotent stem cells through a ∼4 month chimeric culture model in xenogeneic reconstituted ovaries (xrOvaries).^14^ This remarkable work demonstrated that advanced human germ cell types can be differentiated from stem cells *in vitro*, a key step towards IVG. However, this impressive method remains challenging to utilize, particularly in screening paradigms that require scaled, parallelizable cell type differentiation, due to the extensive culture periods in 3D chimeric aggregates and air-liquid interface culture, which require fetal mouse gonad dissection combined with human primordial germ cell-like cell specification. To date, no other method yet exists for the direct generation of DDX4+ germ cells from stem cells in humans.

In mouse models, a number of studies have demonstrated complete female IVG, yielding developmentally competent metaphase II (MII) oocytes from stem cells.^15,16^ Impressively, these various mouse models generated healthy, fertile offspring after fertilization and implantation, showing the power of IVG to successfully model reproductive development from stem cells. In addition, recent mouse studies have demonstrated the utility of transcription factor (TF) based methods for driving and enhancing germ cell development.^17,18^ These TF-based methods benefit from their ease of use, rapid developmental timescales, and high efficiency and can serve as complementary or supplementary differentiation platforms to growth factor only methods. TF overexpression methods in human models have been demonstrated in hPGCLCs, with seminal work identifying that the TFs *SOX17*, *TFAP2C*, and *PRDM1* and the GATA members *GATA3/2* comprise an essential TF network for hPGCLC specification.^1,6,10,11,19^ Overall however, few TF-based human germ cell specification studies have been completed, likely due to a combination of challenging cell culture techniques, difficulty in utilizing tunable genetic engineering methods, and lack of compelling TF target prediction tools. Outside of this upstream TF network, few studies have identified and validated in human models the role of downstream TFs in driving or modulating germ cell development. To date, no TF induction-based methods have been demonstrated for deriving more advanced human germ cell types such as DDX4+ germ cells, oogonia, spermatogonia or oocytes from stem cells.

We hypothesized that TFs that perform highly connected regulatory roles in the final stages of female gamete development may be capable of driving robust germ cell formation and differentiation when overexpressed in stem cells. In this study, we perform a broad TF overexpression screen to assess the effect of overexpressing putative oogenesis-regulating TFs in human induced pluripotent stem cells (hiPSCs) and assay for formation of reporter positive human cell types. Using graph theory algorithms to identify central regulators of oogenesis, we infer TFs that may perform highly connected roles in the regulation of human primordial oocyte to ovulated oocyte development stages.^20^ We then leverage high throughput, monolayer differentiation methods amenable to single factor and combinatorial genetic screening for identifying TFs that increase germ cell reporter-positive cell yield. We identify the TFs, *DLX5*, *HHEX*, and *FIGLA,* whose individual overexpression drives potent enhancement of canonical hPGCLC formation. We likewise identify *LHX8*, *SOHLH1*, and *ZNF281* as TFs whose combinatorial overexpression directly generates DDX4+ induced oogonia-like cell (iOLC) formation in four days in monolayer culture. We furthermore demonstrate a method for feeder-free post-isolation expansion of the DDX4+ iOLCs that utilizes recently described long term hPGCLC expansion and maintenance methods.^7^ Finally, we describe an improved protocol for DDX4+ iOLC production that uses DNA methyltransferase inhibition and overexpression of the additional TFs *ANHX* and *ZGLP1* to achieve higher iOLC yields and an epigenetic state that more closely resembles *in vivo* oogonia. Using immunofluorescence, transcriptomic, and epigenomic assays, we demonstrate these TF-based induced oogonia-like and primordial germ cell-like cells closely resemble *in vivo* and *in vitro* human germ cells. Together, these results build on the foundational platform for targeted derivation of human germ cell types from pluripotent stem cells via TF-directed differentiation. Furthermore, these methods suggest a novel future platform for human IVG, expanding the breadth of genetic engineering and cell-based tools available for reproductive development studies.

## Results

### *In silico* identification of transcription factors that are central regulators of human oogenesis

Previous studies to identify regulators of mouse oogenesis leveraged differential gene expression of the primordial-to-primary follicle transition, identifying TFs that regulated mouse oocyte maturation and were functionally capable of driving oocyte-like cell formation from stem cells.^9^ Similarly, to identify candidate TFs that govern the dynamic gene regulatory networks (GRNs) of human oogenesis, which can thus be prioritized for overexpression studies, we curated publicly-available single cell RNA sequencing (scRNA-seq) data of natural folliculogenesis and normalized the gene expression of germ cell transcriptomes at various stages of follicle development, including primordial, primary, secondary, antral, and preovulatory follicles.^7,14,21,22^ After constructing this pseudo-time series transcriptomic matrix, we employed unsupervised, regression-based GRN inference via regularized stochastic gradient boosting to generate a representative, directed graph of oocyte state during folliculogenesis, where nodes represent genes and edge weights correlate to the regulatory effect between TFs and their downstream targets.^23^ After conducting standard pruning of the graph for low-weight, spurious edges, we applied the adapted PageRank algorithm from our recently-published STAMPScreen pipeline to prioritize the 36 most critical, “central” TFs within the GRN (Figure 1A, Table S1).^20^ Our ranked list includes both previously-identified regulators of mammalian oocyte transcriptional networks and germ cell development, such as *NOBOX*, *LHX8*, and *OTX2*, as well as potential novel regulators, including *ZNF281*, *SOX13*, and *DLX5*.^18,24^ We additionally include 10 TFs that ranked slightly lower on our predictive list (which only leveraged late stage oogenesis data), but are known to perform important roles in earlier specific stages of germ cell development such as PGC formation or that have been identified in similar mouse studies.^10,11,17,18^ The list of the 36 TFs and 10 control TFs is presented in Table S2 with the isoform chosen for screening, if it was predicted or selected as a control, and any known involvement in germ cell development. Associated GO term analysis of the 46 factors is shown in Figure 1B, with reproduction and oogenesis terms highlighted as well as select general GO terms. We furthermore assessed the expression of all 46 TFs in single cell RNA-sequencing (scRNA-Seq) datasets from the recently described human fetal gonad atlas, which is shown in Figure S1.^25^ As can be seen, each of the 46 TFs is expressed in the fetal gonad germ cells in both male and female development. Overall, these results motivated us to screen prioritized candidates in hiPSCs to resolve the human oocyte transcriptional network experimentally.

**Figure 1.**
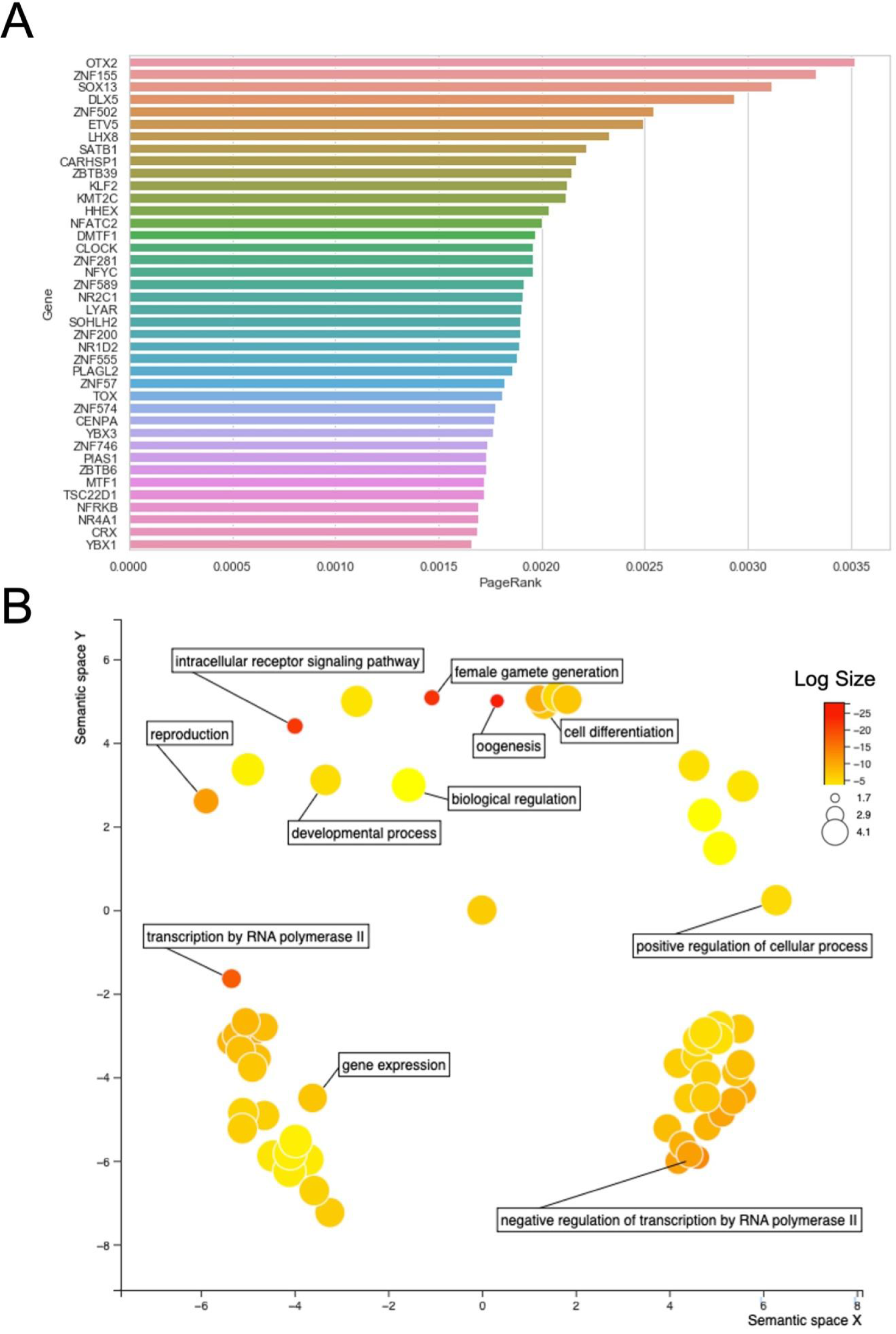
*In silico* identification of TFs with putative germ cell formation regulatory functions. A) Inferred gene regulatory network generated from folliculogenesis dataset of Zhang et al. 2018. with prioritized TF screening list, ordered by PageRank score imputed from the gene regulatory network and centrality algorithm. All 46 TFs utilized in the study were analyzed using the GO term tool g:profiler, and significant GO terms were plotted in Revigo visualizer highlighted terms related to significantly enriched pathways for germline development and transcription factor function. Color and dot size reflect log size of the p-adjusted value.

### Development of a TF-overexpression screening platform in hiPSCs for germ cell-reporter formation

We constructed a <u>D</u>DX<u>4</u>-T2A-td<u>T</u>omato; <u>N</u>ANOS<u>3</u>-T2A-m<u>V</u>enus dual reporter female hiPSC line (D4TN3V) using CRISPR-Cas9-mediated homology directed repair (HDR) (Figure S2A) and confirmed proper reporter insertion by whole genome sequencing (Figure S2B). The cell line was confirmed karyotypically normal and pluripotent using the Karyostat and Pluritest assays (Figure S2C, S2D). Using flow cytometry after CRISPR-activation induction of the DDX4 reporter locus and monolayer hPGCLC induction, we confirmed that the reporter lines induce expression of the fluorescent markers upon induction of the target genes and for the NANOS3 reporter, overlaps with known hPGCLC markers EpCAM/ITGA6 (Figure S2E, S2F). For screening purposes, a four day monolayer protocol inducing through epiblast-like intermediates followed by BMP4 induction was adapted from monolayer hPGCLC generation methods described previously (Figure 2A).^9^ Monolayer methods were chosen for screening due to the ease and scalability of use compared to floating aggregate methods. We generated doxycycline-inducible vectors expressing a full length cDNA for each of the 46 TFs using a highly optimized piggyBac backbone vector for tunable expression in hiPSCs from our STAMPScreen pipeline.^26,27^ We generated 46 individual D4TN3V hiPSC lines harboring integrations of each TF individually through super piggyBac transposase-mediated insertion (Figure S3A).^6,14,28^ Polyclonal pools for each TF were utilized for screening purposes. Through induction of each individual line via doxycycline followed by 3’ UTR barcode capture in RNA-seq, we demonstrated expression of the intended TF and its associated barcode in all 46 lines utilized in the study, confirming each line’s genotype and functional gene expression of each vector (Figure S3B). We additionally utilized the *DLX5* cell line and validated that TF expression in our system is highly linked to doxycycline concentration, as expected, reaching levels of ∼1,000 fold induction compared to the no doxycycline condition in hiPSCs determined by qPCR (Figure S3C).

**Figure 2.**
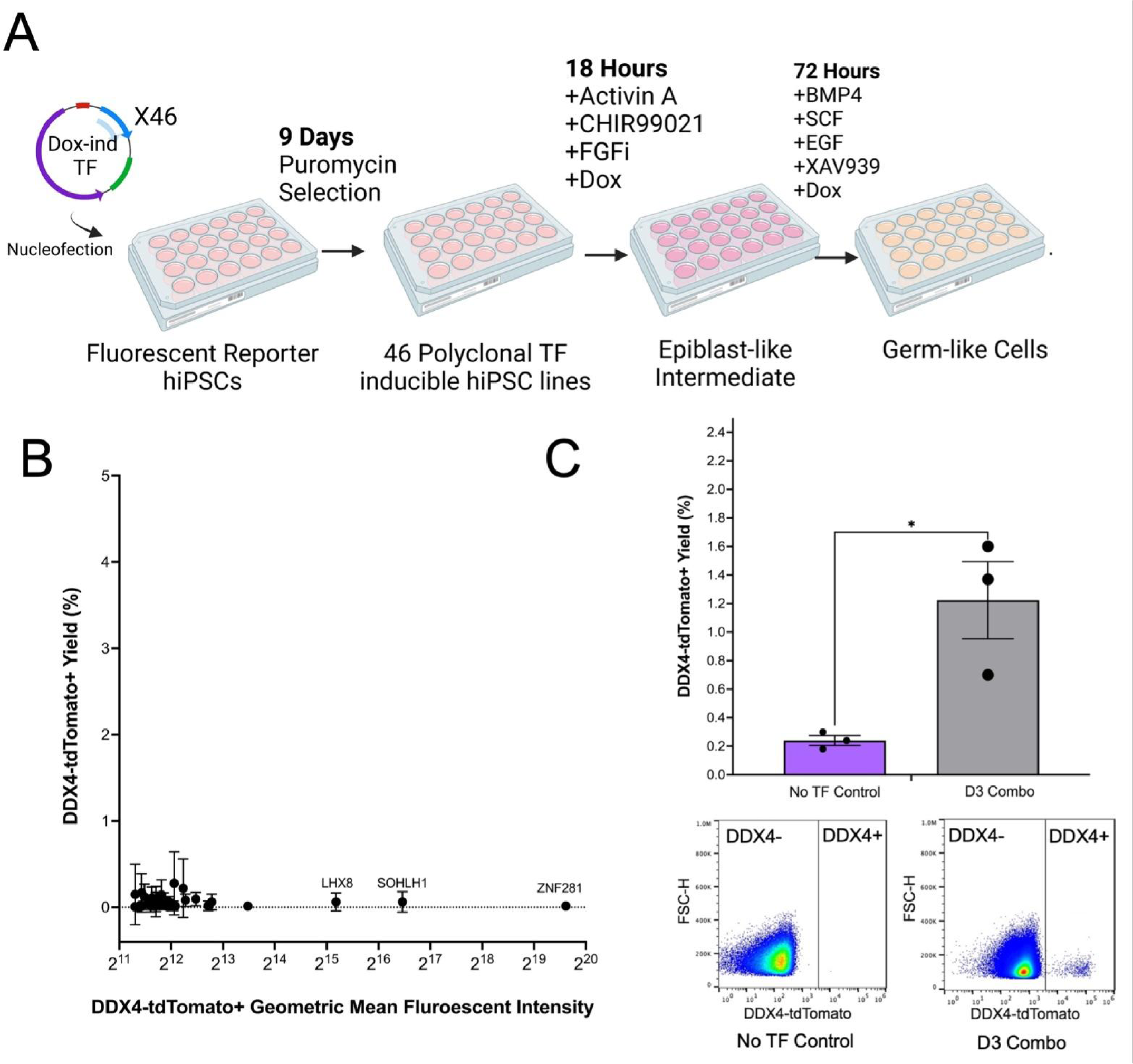
Combinatorial *ZNF281*, *LHX8*, and *SOHLH1* (D3) overexpression drives rapid DDX4+ cell formation. A) Schematic workflow for TF overexpression screening, with time intervals and media supplements. B) Flow Cytometry results of the screen for the DDX4-T2A-tdTomato reporter are visualized with percent reporter positive on the y-axis, and geometric mean fluorescence intensity of the tdTomato+ cells on the y-axis. Error bars represent the SEM of the induction yield between the triplicates in the plus dox condition. Flow cytometry results are visualized for DDX4-T2A-tdTomato reporter yield (percent) for the no TF control and the D3 combo in n=6 replicates (3 independent hiPSC lines, with 2 replicates per line). Individual dots represent the average for the technical replicates for each biological replicate. Data is presented as a mean +/-SEM. Statistical significance was determined by two-sided T-test comparison between each combination and the no TF control, with a p-value <0.05 considered as significant. *** p<0.001, ** P<-.01, * p<0.05. Representative flow cytometry results are visualized for both the no TF control and the D3 combination.

### Identification of TFs regulating reporter positive cell formation via overexpression screening

We first assessed production of DDX4+ cells from hiPSCs via flow cytometry in the presence or absence of doxycycline for all 46 TFs in triplicate (Figure 2A). *DDX4* is broadly expressed throughout germ cell development, arising in gonadal hPGCs after migration to the fetal gonad and increasing in expression during oogonia/spermatogonia formation. Compared to control, the overall percentage of DDX4+ cells was not greatly enriched by any single TF (Figure 2B, y-axis). However, a small percentage of cells with significantly elevated DDX4-reporter expression intensity was identified in the *ZNF281*, *LHX8*, and *SOHLH1* induction conditions (Figure 2B, x-axis). We hypothesized that combinatorial expression of these factors may increase DDX4-reporter yield and generate a more reproducible yield of highly DDX4 reporter-positive cells. To investigate this hypothesis, we generated three independent D4TN3V cell lines harboring a polyclonal integration of all three TFs, which we term <u>D</u>DX4 by <u>3</u> TFs (D3), and assessed the induced DDX4+ yield compared to a no TF control in biological triplicate (Figure 2C). We find the D3 combination drives a statistically significant increase in DDX4-reporter cell yield (1.22% +/-0.27% SEM). Remarkably, this high DDX4+ population is obtained in just 4 days in monolayer through direct TF induction during a primordial germ cell-like cell differentiation protocol. Based on DDX4 expression and subsequent analyses showing similarities between our DDX4+ cells and oogonia, we term these induced oogonia-like cells (iOLCs).

We likewise assessed NANOS3+ cell yield via flow cytometry in the presence or absence of doxycycline for all 46 TFs in triplicate (Figure S4). We show that 23 TFs (15 computationally predicted and 8 controls) drove a statistically significant increase in NANOS3+ cell yield compared to the no TF control, while the other 23 TFs demonstrated a trend towards upregulation, but not significantly. Remarkably, 3 TFs (*DLX5*, *HHEX*, and *FIGLA*) induced a NANOS3+ yield higher (23.6% +/-2.4% SEM, 10.5% +/-0.9% SEM, and 10.3% +/-0.7% SEM) than that of three known TF regulators of hPGCLC development: *SOX17*, *TFAP2C*, and *PRDM1 (8.6% +/-0.5% SEM, 8.5% +/-0.9% SEM, and 8.0% +/-1% SEM)* respectively.^1,11,19^

### TF overexpression drives formation of NANOS3+ cells with canonical hPGCLC characteristics

Utilizing conventional floating aggregate hPGCLC induction methods, we demonstrated that individual overexpression of *DLX5*, *HHEX*, and *FIGLA* drove a statistically significant increase in putative hPGCLC formation as assessed by the canonical surface markers EpCAM/ITGA6 in the plus doxycycline condition (14.6% ± 0.35%, 26.8% ± 0.08%, 26.5% ± 0.46%) versus minus doxycycline condition (3.6% ± 0.24%, 5.4% ± 0.52%, 5.1% ± 0.89%) (Figure S5A).^19^ Putative hPGCLC yield was significantly lower in the no TF control condition and did not significantly differ between the plus doxycycline (4.3% ± 0.63%) and minus doxycycline conditions (4.2% ± 0.20%). We demonstrated through RNA-sequencing that these TF-driven NANOS3+ cells displayed canonical upregulation of *SOX17*, *PRDM1*, *TFAP2C* and downregulation of *SOX2* gene expression compared to the no doxycycline hiPSC controls (Figure S5B). Likewise, using immunofluorescence imaging we demonstrated the unsorted germ cell populations in floating aggregate displayed SOX17, OCT4, ITGA6 triple positive cells, key features of hPGCLCs, which were more abundant in the TF overexpression floating aggregates than in the No TF control, in line with the flow cytometry findings (Figure S6). Together, these results identify individual overexpression of *DLX5*, *HHEX*, and *FIGLA* improves canonical hPGCLC formation.

### iOLCs are capable of post-isolation, feeder-free identity maintenance and expansion and display endogenous DDX4 protein expression

It is known that *in vivo*, primordial germ cells and oogonia develop and mitotically expand, establishing the human ovarian reserve.^14,29^ Recent methods have been developed for expansion of hPGCLCs *in vitro* post-isolation.^5,7,30^ No method yet exists for the maintenance of human oogonia or oogonia-like cells *in vitro* after FACS isolation. We thus sought to determine if our D3 iOLCs could be maintained in culture post-isolation. Serendipitously, we found that by seeding isolated iOLCs onto matrigel-coated plates feeder-free in a media composition recently described for hPGCLC expansion, we were able to maintain and expand the DDX4+ cell population for at least 28 days (Figure 3A).^7^ After FACS isolation, these D3 DDX4+ cells slowly divided for 28 days, retaining expression of DDX4, as can be seen by live cell fluorescent imaging of the tdTomato reporter protein. These DDX4+ retained robust expression of the DDX4-T2A-tdTomato reporter, with clear visualization of the cleaved, diffuse fluorescent protein. Since our reporter was not a fusion protein to DDX4 itself, we saw the expected pattern of fluorescent protein in both the nucleus and cytoplasm as opposed to just the cytoplasm. Additionally, we performed immunofluorescence imaging on fixed cells from the sorted DDX4+ cells at day four and day 28 of *in vitro* expansion and found they retained expression of endogenous DDX4 and only weak expression of DAZL and no expression of OCT4 (Figure 3B). Intriguingly, we found that at day 28 the majority of the DDX4+ cells existed in a multi-nucleated state reminiscent of the cell state of oogonia found *in vivo* in germ cell nests.^31^ These multi-nucleated cells showed a single cytoplasm and 4-8 nuclei per cell and were DDX4+, OCT4- and weakly DAZL+ (Figure 3B). We therefore conclude that our TF-derived iOLCs are capable of feeder-free maintenance and expansion of DDX4+ identity post-isolation in a multi-nucleated state.

**Figure 3.**
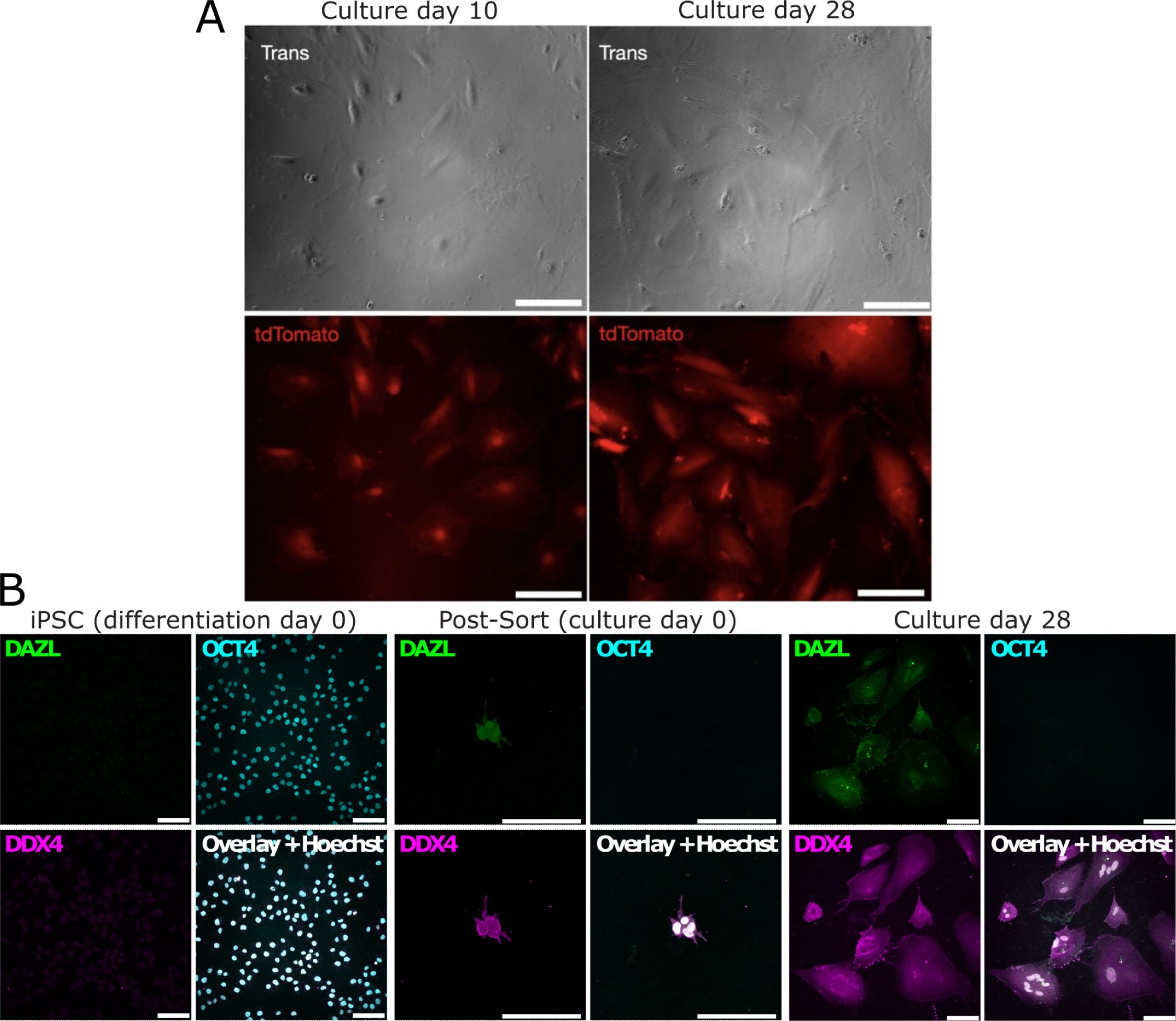
D3 iOLCs are capable of post-isolation feeder-free maintenance and expansion and display continued DDX4 protein expression. A) Live cell fluorescent imaging is performed on FACS isolated D3 combination-derived iOLCs grown feeder-free on matrigel coated plates at day 10 and 28 post-FACS isolation. Transmission light microscopy is used to visualize cells (top panel) and tdTomato fluorescent protein derived from the cleaved DDX4-T2A-tdTomato reporter is visualized (bottom panel) using the Echo Revolve fluorescent microscope on the Texas Red setting. Scale bar is 90µm. B) DDX4+ cells from D3 combination were FACS isolated at day 4 of differentiation. Cell fixation and staining was performed at day four and day 28 of post-isolation culture and stained for Hoechst (blue), DAZL (green), and DDX4 (red), and OCT4 (gray). Representative images were taken using confocal microscopy. Scale bar is 100µm.

### An improved protocol allows production of iOLCs with higher efficiency

Although expression of our D3 TF combination reliably produces DDX4+ iOLCs, the efficiency is low. Therefore, we sought to identify factors which could improve this. We generated a library of barcoded PiggyBac transposon plasmids for Dox-inducible expression of 62 additional oogonia-related TFs and RNA-binding proteins, as well as 31 constitutively active signaling proteins.^32^ Additionally, we included a CRISPRi vector targeting *DNMT1*, since suppression of DNA methylation is known to be crucial for germ cell development.^33^ We integrated this library into cells which already contained Dox-inducible D3 TFs, induced oogonia-like cells, and performed barcode enrichment screening (Figure 4A). We observed that the top hit was *DNMT1* CRISPRi, and we selected a sub-library of 24 factors for a follow-up round of barcode enrichment screening (Figure 4B), which we performed in both male and female DDX4 reporter hiPSCs. In this screen, we treated the cells with 5 µM GSK-3484862, a noncovalent inhibitor of DNMT1,^34,35^ to inhibit DNA methylation. Additionally, we switched to a basement membrane extract overlay monolayer induction protocol which allowed higher cell seeding densities.^36^ In some samples, we also tested the retinoic acid receptor agonist AM580, but we observed that this inhibited cell proliferation, and we did not include in in the final protocol out of concerns that it could induce premature differentiation of oogonia. After this second round of screening, we chose six promising factors and performed a fractional factorial screen, testing 16 combinations in two cell lines. These experiments identified *ANHX* and *ZGLP1* as TFs which could increase the yield of DDX4+ iOLCs when co-expressed with the D3 TFs (Figure 4C). We term this improved combination D5.

**Figure 4.**
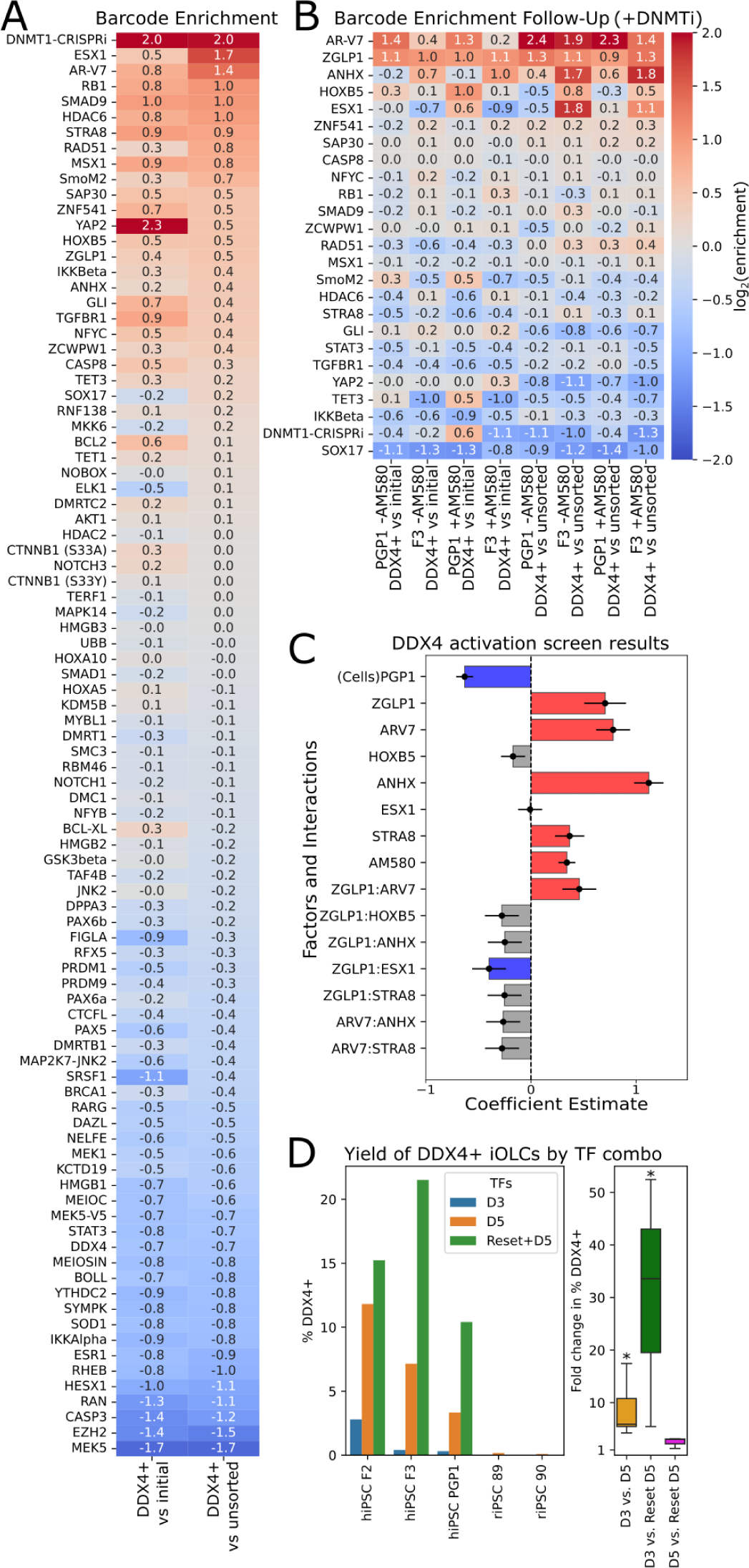
Optimization of the iOLC induction protocol. (A) Initial barcode enrichment screen. Factors are plotted in order of log2-fold enrichment in DDX4+ cells vs. unsorted cells. (B) Followup barcode enrichment screen. A smaller set of factors were tested in an increased number of conditions. +AM580 denotes addition of the retinoic acid receptor agonist AM580. In all conditions for this experiment, cells were treated with DNMT1 inhibitor, meaning that the effect of DNMT1 CRISPRi was masked. (C) Results of a fractional factorial screen of top factors identified in the previous experiment. Sixteen combinations were tested in two cell lines. A linear model was fit to the logit-transformed percentage of DDX4+ cells. Coefficients and standard errors are plotted. (D) Comparing yield of iOLCs with D3 vs. D5 TFs in the improved protocol (BME overlay induction, +DNMT1 inhibitor). A total of three human and two rhesus monkey iPSC lines were tested. Additionally, transient naïve resetting was tested in combination with D5 TFs for three human iPSC lines.

We validated the improved performance of the D5 TFs by comparing them with D3 TFs in three hiPSC DDX4-T2A-tdTomato reporter lines (two female, one male), and two rhesus macaque iPSC DDX4-T2A-tdTomato reporter lines (both male). Among all five lines tested, the yield of DDX4+ cells was significantly higher using D5 TFs (average 8.75-fold increase; paired *t*-test, *p*=0.02) (Figure 4D). We additionally tested using transient naïve resetting to improve the yield of iOLCs, since this had been reported as a method for efficient germ cell induction.^37^ Naïve resetting of hiPSCs with Sox2^A61V^-Klf4-Myc episome,^38^ followed by iOLC induction using D5 TFs, allowed the generation of DDX4+ cells with good efficiency (Figure 4D). The yield was significantly higher than for D3 TFs (average 30.5-fold increase; paired *t*-test, *p*=0.03) but the increase in yield over D5 TFs did not reach statistical significance (average 2.5-fold increase; paired *t*-test, *p*=0.10). Overall, this optimized protocol, which combines additional TFs, DNMT1 inhibition, and higher cell seeding density, enabled us to access larger numbers of iOLCs for use in subsequent experiments.

### iOLCs generated using the improved protocol show robust DNA methylation erasure similar to *in vivo* oogonia

During germline development, cells proliferate in the absence of maintenance DNA methyltransferase activity, leading to genome-wide erasure of DNA cytosine methylation.^33^ This is a crucial process for mammalian gametogenesis, as it erases parental imprints and allows the establishment of sex-specific imprinting. Furthermore, crucial genes for germ cell development, such as *DAZL*, are expressed from methylation-sensitive promoters.^39,40^ Notably, previous TF-driven methods for generating oocyte-like cells from mouse pluripotent stem cells did not achieve DNA methylation erasure nor expression of *Dazl*.^18^ However, previous xrOvary based methods for generating human oogonia-like cells were able to erase DNA methylation over a culture period of 77–120 days.^14^

Given the shorter culture period (4–5 days) of our method, we investigated whether our iOLCs were also erasing their methylation. We performed whole genome enzymatic methyl sequencing^41^ on a total of 18 samples, including primed hiPSCs, transient naïve reset hiPSCs, hPGCLCs, and iOLCs generated using our original and improved protocols. Notably, the improved protocol included a chemical inhibitor of maintenance DNA methyltransferase DNMT1. We first analyzed overall methylation levels (Figure 5A), comparing our samples to previously published *in vivo* hPGC data as well as data from the xrOvary culture system.^14,42^ We observed that iOLCs from the original protocol, which did not include a DNMT1 inhibitor, did not have lower DNA methylation relative to hiPSCs. hPGCLCs showed a slight decrease in methylation. Encouragingly, iOLCs from the new protocol were strongly demethylated, reaching levels similar to xrOvary and hPGCs.

**Figure 5.**
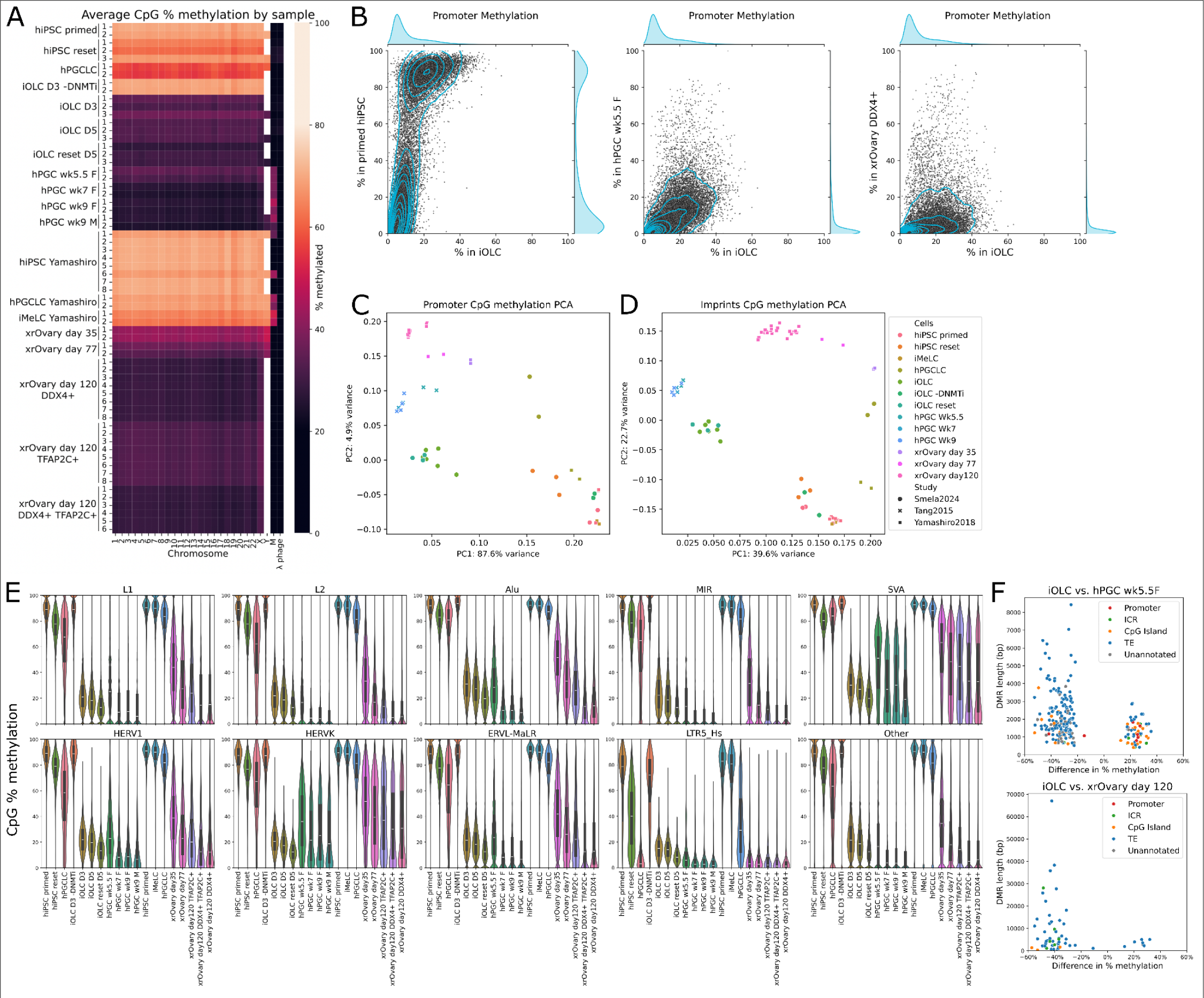
Epigenetic analysis of iOLCs in comparison to hPGCs and xrOvary DDX4+ cells. (A) Heatmap showing average % CpG methylation for each chromosome in each sample. (B) Comparing average methylation for each promoter in iOLCs vs. primed hiPSCs, hPGCs (female week 5.5), and xrOvary day 120 DDX4+ cells. Promoter methylation is efficiently erased in iOLCs. (C) Principal component analysis for promoter methylation among different sample types. Along PC1 (which captures 87.6% of variance), iOLCs are similar to hPGCs and xrOvary cells. (D) Principal component analysis for imprint methylation among different sample types. iOLCs are most similar to hPGCs. (E) Distribution of methylation for different classes of TEs among different sample types. (F) Differentially methylated regions between iOLCs and hPGCs, and between iOLCs and xrOvary day 120 DDX4+ cells. Regions are colored based on annotated features. Most correspond to TEs.

We next examined DNA methylation at functionally relevant loci including promoters (Figure 5B/C), CpG islands (Figure S7A), imprinting control regions (ICRs) (Figure 5D), and transposable elements (TEs) (Figure 5E). In hiPSCs, promoters and CpG islands displayed bimodal methylation (Figure 5B and S7A). Most (56% of promoters and 61% of CpG islands) had a low (0-20%) level of methylation, and a smaller fraction (24% of promoters and 29% of CpG islands) had a high level (80-100%). By contrast, iOLCs had no promoters or CpG islands above 80% methylation, and the proportion with lower than 20% methylation was greatly increased (73% - 92%). Notably, the methylation of the *DAZL* promoter decreased from 96% in hiPSCs to 26% in iOLCs. To evaluate the epigenetic similarity of our iOLCs with xrOvary cells and hPGCs, we performed a principal component analysis of promoter methylation. Along principal component 1, which captured most of the variance, our iOLCs were similar to xrOvary oogonia and hPGCs (Figure 5C).

Erasure of parental methylation at ICRs is crucial for the generation of functional gametes.^33^ We examined DNA methylation at 332 candidate human ICRs found by a previous study,^43^ which identified regions with 50±15% methylation in somatic tissues and 0 or 100% methylation in eggs and sperm. Interestingly, we found that these regions had an average of 76% methylation in our hiPSC samples. Previous studies in mouse and human iPSCs identified aberrant ICR hypermethylation as a consequence of iPSC reprogramming,^44,45^ and this may have also occurred in our hiPSCs. However, in iOLCs, only 12 out of 327 ICRs retained >40% methylation, with the majority having <15% methylation. A principal component analysis showed that our iOLCs were more similar than hiPSCs or xrOvary oogonia to *in vivo* hPGCs (Figure 5D). Therefore, despite the relatively short duration of the iOLC induction protocol, iOLCs largely completed the process of imprint erasure.

Although genome-wide DNA demethylation is necessary for gametogenesis, demethylation of TEs can activate the expression of these potentially deleterious elements. In the human germline, certain TEs retain DNA methylation despite genome-wide demethylation.^42^ This “escape” from demethylation ensures that the TEs remain suppressed. Given that our protocol included a small-molecule inhibitor of DNMT1, we wondered if any TEs would escape demethylation. Examining methylation levels at 3,213,373 TEs, we found that most were demethylated in iOLCs (average methylation 22%) (Figure 5E). However, we identified 280 “escapee” TEs that retained >80% methylation, and 8657 TEs which retained >60% average methylation. Thus, although 5 days of GSK-3484862 treatment was sufficient to erase DNA methylation at promoters and imprinting control regions, a subset of TEs remained methylated. Interestingly, this set of escapee TEs showed little overlap with escapee TEs observed *in vivo* or in xrOvary oogonia (Figure S7B).

Finally, we identified and annotated differentially methylated regions (DMRs) between our iOLCs and xrOvary and fetal oogonia. Among the *in vivo* germ cells,^42^ our iOLCs were most similar to week 5.5 female samples. There were 58 regions significantly more methylated than *in vivo*, and 173 loci significantly less methylated (Figure 5F). Most DMRs were associated with TEs (154 regions), and only a small minority were associated with promoters (8 regions) or imprint control regions (8 regions). Among the xrOvary cells, our iOLCs were most similar to day 120 DDX4-positive cells. There were a total of 7 regions where iOLCs were significantly more methylated than xrOvary cells, and 54 significantly less methylated. Again, most of these DMRs (54 regions) were associated with TEs, and in this case none were associated with promoters. By contrast, iOLCs generated using the protocol without DNMT1 inhibition had 208,715 DMRs, all of which were more methylated. Overall, our data indicate that five days of DNMT1 inhibition effectively resets DNA methylation to an oogonia-like state except at a few regions, most of which are associated with TEs.

### Transcriptomic analysis of iOLCs shows similarities with human fetal germ cells and xrOvary oogonia

We next profiled the gene expression in iOLCs. To investigate the effects of different iOLC induction methods, we induced iOLCs using our original protocol (D3 TFs without DNMT1 inhibition) and improved protocol (with DNMT1 inhibition, and either D3, D3+ZGLP1, D3+ANHX, or D5 TF combinations). Additional samples included iOLC TFs in combination with a constitutively active androgen receptor splice variant (AR-V7), to see if this would upregulate spermatogonia-related genes. We harvested iOLCs from each condition and performed bulk RNA-seq on the sorted DDX4+ cells. For the D5 TF combination, we also sequenced RNA from the unsorted cell population.

First, we examined the expression of known marker genes for different stages of germline development: pluripotent stem cells, primordial germ cells, oogonia, meiotic germ cells, and oocytes (Figure 6, Table S3). As a control, we used previously published data from human fetal germ cells and xrOvary oogonia.^14,42,46^ We found that iOLCs from all induction protocols expressed the oogonia marker genes *DDX4, DAZL, STK31,* and *MAEL.* However, iOLCs from the original protocol retained expression of the pluripotency/PGC marker *POU5F1,* and lacked expression of other oogonia markers such as *MAGEB1/2, MSX1, TKTL1,* and *OOEP*. By contrast, iOLCs from the improved protocol expressed these oogonia markers and had lower expression of *POU5F1*, similarly to xrOvary day 120 DDX4+ TFAP2C-oogonia. Additionally, iOLCs upregulated some meiotic markers, including *CTCFL, SYCP1/2/3, HORMAD1,* and *TEX11/12*. However, these cells were likely not fully committed to meiosis, as they had lower expression of other markers such as *REC8* and *SPO11*. Similarly, iOLCs did not express genes related to oocyte growth, such as *NOBOX, GDF9, BMP15, ZP3* and *NPM2*.

**Figure 6.**
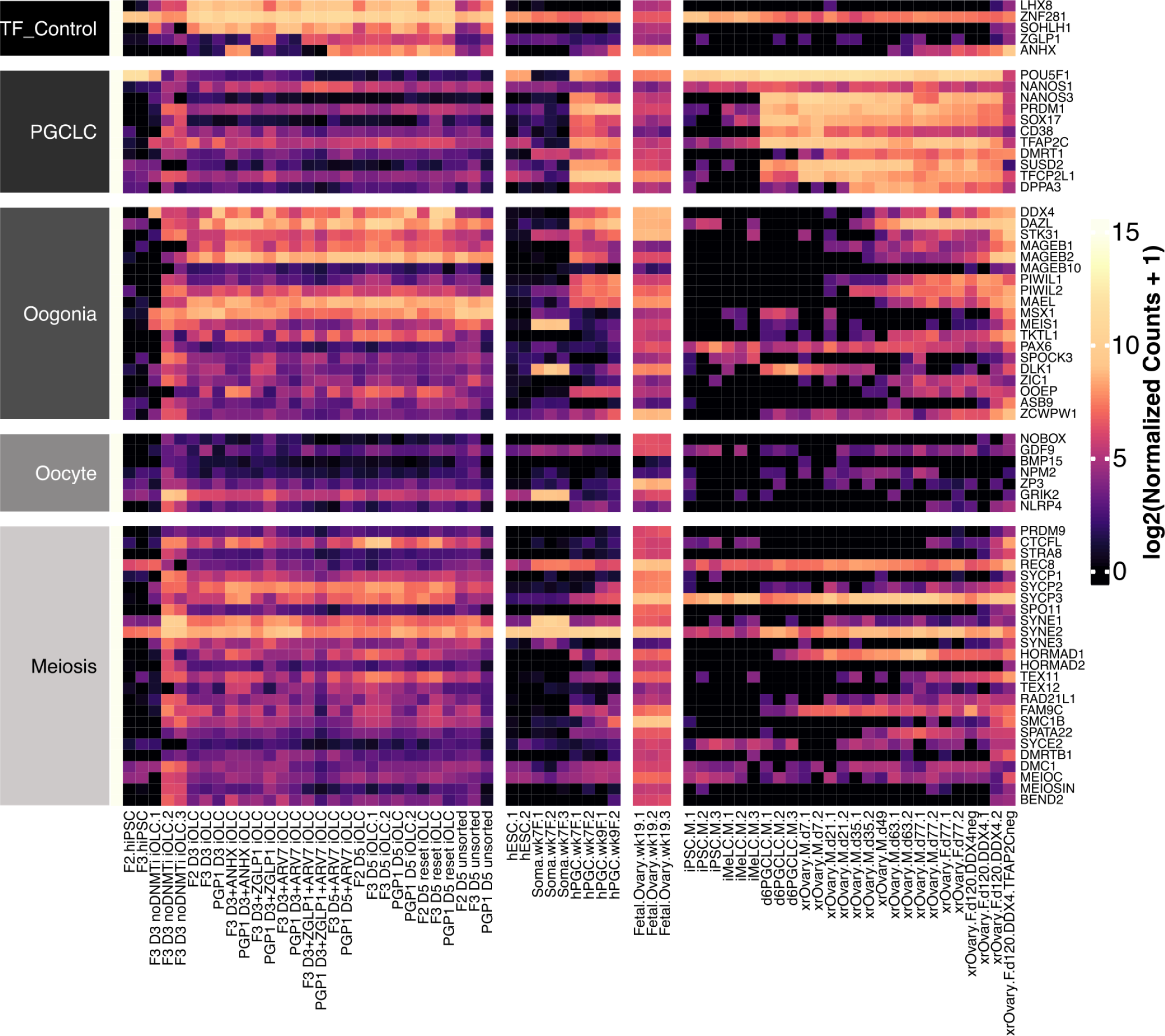
Heatmap of marker gene expression. From left to right, samples include: hiPSCs, iOLCs generated with D3 TFs and without DNMT1 inhibition, iOLCs generated with various TF combinations using the improved protocol, unsorted cells from iOLC induction; Tang *et al*. hESCs, fetal ovarian somatic cells, and female hPGCs; Yatsenko *et al*. week 19 fetal ovarian germ cells; and Yamashiro *et al*. hiPSCs, hiMeLCs, hPGCLCs, and xrOvary germ cells ordered by developmental stage.

Next, we performed a differential gene expression analysis to identify differences among iOLCs induced using different TFs. In comparison to the D3 TF combinations, the other TF combinations showed significant changes to gene expression (Table S4). We conducted a GO term enrichment analysis on differentially expressed genes. AR-V7 overexpression did not result in activation of spermatogenesis-related genes; the only enriched GO term among upregulated genes was “chromatin organization”, and contrary to our expectations, “male sex differentiation” was enriched among downregulated genes. We found that overexpression of ANHX (relative to D3 control) upregulated genes related to reproductive development (including “spermatogenesis”, “sexual reproduction”, “gamete generation”, “meiotic cell cycle”, “developmental process involved in reproduction” and “retrotransposon silencing”), and downregulated genes related to mesenchymal and cardiac development. For overexpression of ZGLP1, no GO terms were significantly enriched in the differentially expressed genes. However, the D5 combination (overexpression of both ANHX and ZGLP1) showed more extensive changes. Upregulated genes were enriched for GO terms related to metabolism, meiosis, germ cell development, and oogenesis. (Table S5). GO terms enriched among downregulated genes were mainly related to cell adhesion, migration, blood vessel development, and angiogenesis. These results indicate that the two additional factors in the D5 combination act to reinforce germ cell development and downregulate off-target differentiation.

We additionally compared gene expression in D5 DDX4+ iOLCs with unsorted cells (largely DDX4-) from the same induction. Relative to the iOLCs, the bulk population expressed less *DDX4* (as expected), and also had lower expression of other oogonia markers *DAZL, BOLL, TEX15* and *STK31*. Significantly downregulated GO terms were related to gametogenesis, meiosis, and metabolism (Table S5). GO terms upregulated in the bulk population relative to iOLCs were related to vascular, cardiac, and nervous development. We also compared gene expression in D5 DDX4+ iOLCs generated from primed hiPSCs vs. transient naïve reset hiPSCs. There were only a few significant differentially expressed genes (one downregulated and seven upregulated). However, we note that both *SOX17* and *GATA4* were significantly upregulated in iOLCs derived from reset hiPSCs, and these transcription factors are known to be expressed in fetal germ cells.^6,47^

Finally, we examined differences between our iOLCs and *in vivo* and xrOvary germ cells. Unfortunately, our efforts at transcriptome-wide comparison between studies were hindered by batch effects, likely due to different sequencing techniques. This was clearly shown by a principal component analysis in which pluripotent stem cells from different studies (our hiPSCs, Yamashiro *et al.* hiPSCs, and Tang *et al.* hESCs) did not cluster together (Figure S8). However, we took advantage of the fact that a known cell type (pluripotent stem cells) was sequenced in each of these studies to perform batch correction by adding the study as a covariate to our differential expression model.

We performed a differential gene expression and GO term enrichment analysis between our D5 iOLCs and hPGCs and xrOvary day 120 oogonia. As a control, we also compared our iOLCs with fetal ovarian somatic cells from the Tang *et al*. dataset, and found that GO terms related to germ cells and meiosis (e.g. “meiotic cell cycle”, “sexual reproduction”, and “gamete generation”) were significantly enriched among genes that were less expressed in fetal somatic cells relative to iOLCs. By contrast, we did not observe significant enrichment of GO terms related to germ cell development between iOLCs and hPGCs or xrOvary oogonia. Differentially expressed genes were mainly associated with metabolic processes such as “nitrogen compound metabolic process”, or generic terms such as “multicellular organismal process” (Table S5). However, terms related to meiosis, such as “negative regulation of mitotic cell cycle”, “meiotic cell cycle”, “DNA damage response”, and “double-strand break repair via homologous recombination” were upregulated in xrOvary day 120 oogonia relative to our iOLCs. This may reflect that xrOvary day 120 oogonia are beginning meiosis, whereas our iOLCs are at an earlier developmental stage. Although these results should be incorporated with caution given the possibility of incomplete batch correction, it appears that our iOLCs express broadly similar levels of germ cell related genes compared with hPGCs or xrOvary oogonia.

## Discussion

In this study, we identified 36 TFs that may play regulatory roles within the underlying GRNs of gametogenesis and further screened these TFs alongside known control TFs via combinatorial and individual overexpression in a germline-reporter hiPSC line. We find three TFs, *ZNF281*, *LHX8*, and *SOHLH1*, drive DDX4+ induced oogonia-like cell formation, while three other TFs, *DLX5*, *HHEX*, and *FIGLA*, individually drive enhancement of hPGCLC yield when overexpressed during hPGCLC specification.

We demonstrate that overexpression of *DLX5*, *HHEX*, and *FIGLA* drive on-target canonical hPGCLC formation, shown in the expression of the core primordial germ cell genes *SOX17*, *TFAP2C*, and *PRDM1* and proteins SOX17, OCT4, ITGA6, and EpCAM in floating aggregate differentiation methods. While the overall hPGCLC yield, particularly in the no TF controls, was generally low with the D4TN3V reporter line utilized in this study, this yield was in line with the reported yields seen in other studies which have noted a wide variety in absolute hPGCLC yield between different hiPSC lines.^12^ Interestingly, overexpression of *DLX5*, *HHEX*, and *FIGLA* improved the hPGCLC yield to a level that is similar to high performing hiPSC lines noted in other studies, and may represent a novel method for improving hPGCLC yield in difficult, low yielding cell lines.^6,12,14,19^ More studies will be needed to determine if these three TFs perform germ cell regulating roles *in vivo* in addition to serving as useful cell engineering tools in our described context. It is unclear from our system whether this is the canonical function of the TFs normally during hPGCLC formation or a result of forced overexpression. For *FIGLA*, in particular, it is likely that our overexpression drives a non-canonical function that is not normally seen during hPGC formation, as *FIGLA* is not normally expressed at PGC developmental timepoints *in vivo*.^22^ Nonetheless, this study motivates utilization of our toolkit in a diverse range of hPGCLC screening modalities, which may uncover the role of new TFs in regulating gametogenesis and further elucidate the underlying gene regulatory network of primordial germ cell specification.

We show that overexpression in combination of *SOHLH1*, *LHX8*, and *ZNF281* (D3) drives formation of DDX4+ iOLC formation in just four days. Our DDX4+ cell formation is rapid compared to existing methods which generally require ∼70 to 120 days in xrOvary based methods compared to our four day protocol.^14^ In addition, our DDX4+ cells are generated feeder-free using a 2D monolayer culture protocol, making scaling for production and screening simple and amenable to integration with current high throughput screening designs. Additional studies will be needed to determine the *in vivo* role of these TFs in oogonia and oocyte formation and whether our overexpression-based method for generating these cell types recapitulates the endogenous genetic regulation of oogenesis. As *ZNF281*, *LHX8*, and *SOHLH1* are highly expressed during human folliculogenesis, linked as causal genetic determinants of infertility when mutated, and in the case of, *Lhx8* and *Sohlh1*, found in mouse models to induce oocyte-like formation from stem cells, it is highly likely they play critical roles during *in vivo* oogenesis.^18,21,48^ Expression of the additional TFs *ANHX* and *ZGLP1* (in the D5 combination) increases the yield of iOLCs, upregulates genes related to oogenesis, and downregulates genes related to off-target cell types. *ANHX* is known to be expressed in oogonia,^25^ but its functional role has not yet been previously characterized. In mouse germ cells, *Zglp1* overexpression is known to induce meiosis,^17^ but in human germ cells *ZGLP1* is expressed in oogonia that are not yet activating meiosis.^25^ Via transcriptomic analysis, we find that our TFs drive formation of DDX4+ cells that exhibit upregulation of key oogonia genes, which we designate as induced oogonia-like cells. We find our iOLCs broadly share similarity with *in vivo* fetal oogonia.^21,22^ We also demonstrate maintenance feeder-free post-isolation of our iOLCs using media developed for hPGCLC maintenance, potentially expanding the utility of this cell type for long term studies.^7^

In our optimized protocol for iOLC production, we found that inhibition of DNMT1 efficiently erases DNA methylation and leads to an epigenetic state closely resembling that of human fetal germ cells. We took advantage of a recently described noncovalent DNMT1 inhibitor,^34,35^ which lacks the toxicity of previous inhibitors such as 5-azacytidine. DNMT1 inhibition results in genome-wide demethylation through lack of maintenance methylation during cell divisions, similar to the process that takes place in the human germline.^33^ In mouse germ cells, Dnmt1 is largely blocked from maintaining genome-wide methylation, but retains activity at specific loci related to transposable elements, imprinting control regions, and promoters of meiotic genes.^49^ These loci are actively demethylated by Tet enzymes prior to meiotic initiation. Notably, conditional knockout of Dnmt1 in mouse PGCs results in precocious expression of *Dazl* and differentiation to oocytes and prospermatogonia.^49^ Chemical inhibition of DNMT1 in our protocol may have similar effects. Although we did observe some TEs escaping DNA demethylation in our iOLCs, it is possible that this reflects *de novo* methylation by DNMT3. If DNMT1 retains activity *in vivo* at TE-associated loci, this could explain the TE-associated differential methylation we observed between *in vivo* hPGCs and our iOLCs.

More research is needed to determine the functionality of these iOLCs and whether they can contribute to further gametogenesis and oocyte formation, such as by entering and performing meiosis and oocyte maturation. Additional research as well combining our iOLC methods with co-culture methods such as xrOvaries,^50^ or newer all-human methods such as those described recently,^51^ are needed to determine if these DDX4+ cells are capable of meiotic entry or oocyte maturation. Nonetheless, our rapid and direct generation of human iOLCs provides a novel, reproducible, and simple-to-obtain source of important germline models that can be used in reproductive development and genetic screening studies as models of inaccessible *in vivo* counterparts. In conclusion, we identify novel TFs in driving germ cell formation, establishing a rapid, high-throughput platform for *in vitro* human gametogenesis and reproductive modeling in a simple and efficient manner.

## Methods

### Ethics Statement

All experiments on the use of hiPSCs for the generation of hPGCLCs and oogonia-like cells were approved by the Embryonic Stem Cell Research Oversight (ESCRO) Committee at Harvard University.

### Gene Regulatory Network Inference and TF Prioritization

RNA-seq datasets were obtained from the Gene Expression Omnibus (GEO) database, and log2fc values for each aligned gene for each sample were calculated using the DESeq2 package.^52^ Gene regulatory networks were inferred utilizing the GRNBoost2 algorithm in the Arboreto computational framework.^23,52^ PageRank was calculated for each transcription factor in the resulting network via the NetworkX package, and ranked factors were visualized using Seaborn. The full code and corresponding Jupyter notebooks for the standard STAMPScreen pipeline can be found at: https://github.com/programmablebio/stampscreen.^20^ Gene Ontology Analysis was performed by input of the prioritized TF screening list to the g:profiler tool and selecting the significantly enriched BP terms. g:GOSt performs functional enrichment analysis, also known as over-representation analysis (ORA) or gene set enrichment analysis, on input gene list. It maps genes to known functional information sources and detects statistically significantly enriched terms. These significant GO terms, selected by the adjusted p-value, were then input to the visualizer tool Revigo and plotted as a scatterplot. The axes in the plot have no intrinsic meaning. Revigo uses Multidimensional Scaling (MDS) to reduce the dimensionality of a matrix of the GO terms pairwise semantic similarities. The resulting projection may be highly non linear. The guiding principle is that semantically similar GO terms should remain close together in the plot.

### Cell lines used

For these studies we primarily used the ATCC-BXS0116 female hiPSC line, which we term F3. This cell line was determined to be karyotypically normal via the Karyostat assay and scored as normal in the Pluritest compared to the assays’ control dataset (ThermoFisher). For validation of iOLC production, we also used the ATCC-BXS0115 female hiPSC line (which we term F2), the PGP1 male hiPSC line, and male rhesus iPSC lines 89 and 90.^53^ The rhesus iPSC lines were a gift from Prof. Amander Clark.

### iPSC Culturing

All hiPSCs were maintained feeder-free on hESC-qualified Matrigel coated plates (Corning), as the manufacturer suggested dilution. hiPSCs were maintained in mTeSR media, with mTeSR Plus utilized for standard expansion, passaging and cell line creation and mTeSR1 utilized prior to induction where specified. hiPSCs were passaged in mTeSR1 for at least one passage prior to differentiation in order to remove the stabilized FGF present in mTeSR Plus. Cells were passaged 1:10 to 1:20 every 3-4 days using Accutase and seeded in the presence of 10 µM Y-27632, with media being changed every day for mTeSR1 or every other day when mTeSR plus was utilized. Rhesus iPSCs were cultured on mitomycin-inactivated DR4 MEFs (ATCC #SCRC-1045) in PluriStem medium supplemented with 1 µM XAV939, and passaged using Accutase. Cells were regularly tested for mycoplasma contamination using the ATCC Universal Mycoplasma Detection PCR kit.

### Generation of gametogenesis cell reporter hiPSC lines

Homology arms for target genes (*DDX4, NPM2,* and *NANOS3*) were amplified by PCR from genomic DNA. For each gene, a targeting plasmid, containing an in-frame C-terminal T2A-fluorescent reporter of either *tdTomato* (for *DDX4*), mGreenLantern (for *NPM2*), or *mVenus* (for *NANOS3*), as well as a *Rox-PGK-PuroTK-Rox* selection cassette, was constructed by Gibson assembly. The plasmid backbone additionally had an MC1-DTA marker to select against random integration. sgRNA oligos targeting the C-terminal region of target genes were cloned into the pX330 Cas9/sgRNA expression plasmid (Addgene 42230). For generation of the reporter line, 2 μg donor plasmid and 1 μg Cas9/sgRNA plasmid were co-electroporated into F3 hiPSCs, which were subsequently plated in one well of a 6-well plate. Electroporations were performed using a Lonza Nucleofector with 96-well shuttle, with 200,000 hiPSCs in 20 μL of P3 buffer. Pulse setting CA-137 was used for all electroporations. Selection with the appropriate agent was begun 48 hours after electroporation and continued for 5 days.

After selection with puromycin (400 ng/mL), colonies were picked manually with a pipette. The hiPSC lines generated were genotyped by PCR for the presence of wild-type and reporter alleles. Homozygous clones were further verified by PCR amplification of the entire locus and Sanger sequencing. To excise the selection cassette, hiPSCs were electroporated with a plasmid expressing Dre recombinase. Selection was performed with ganciclovir (4 μM) and colonies were picked as described above. The excision of the selection cassette was verified by genotyping. Reporter lines were screened for common karyotypic abnormalities using a qPCR kit (Stemcell Technologies) followed by verification via Thermo Fisher Cell ID + Karyostat and Pluritest services. Additionally, whole genome sequencing was performed (Novogene, 10X coverage, 150bp paired-end reads) to further verify the reporter alleles. Reads were aligned, analyzed and visualized using the SeqVerify pipeline (version 1.1.0).^54^

### cDNA vector creation

Vectors for cDNA overexpression were generated via MegaGate cloning. Full length cDNAs for each TF of interest were either derived from the human ORFeome or synthesized as full-length constructs. All ORFs were cloned into pENTR221 with stop codons and minimal Kozak sequences. MegaGate was utilized to insert ORFs into the final PB-cT2G-cERP2 3’ UTR barcode-modified expression vectors (Addgene 175503). Three unique barcodes were selected for each ORF with an average hamming distance of six. The three barcoded vectors for each ORF were then pooled, such that for each individual TF there was a mixture of three barcoded vectors. Sanger sequencing was performed across the entire ORF length to confirm canonical sequence with no amino acid changes. For *DNMT1* CRISPRi, the dCas9-KRAB-meCP2 repressor (Addgene 110821)^55^ was used. Three plasmids were constructed with different sgRNAs targeting the promoter of *DNMT1* (GGCTTCAGCAGACGCGGCGG, GGTACGCGCCGGCATCTCGG, GGAGGCTTCAGCAGACGCGG), and these were used as an equal-ratio mixture.

### Generation of inducible TF hiPSC cell lines

Expression plasmids containing TF cDNAs under the control of a doxycycline-inducible promoter were integrated into hiPSCs using piggyBac transposase. To perform the integration, 100 fmol of TF cDNA plasmid, 200 ng piggyBac transposase expression plasmid, and 200,000 hiPSCs were combined in 20 μL of Lonza P3 buffer and electroporated using a Lonza Nucleofector 4D. Pulse setting CM-113 was used for all electroporations. After electroporation, cells were seeded in 24-well plates in mTeSR Plus + 10 μM Y-27632. Selection with 400 ng/ml puromycin began 48 hours after electroporation and continued for 3-5 days. Cells were then passaged without drug selection for 3 days to allow for non-integrated plasmid loss. Finally, cells were again passaged under drug selection to generate a pure, polyclonal integrant pool. Presence and approximate copy number of integrated TF plasmids was confirmed by qPCR on genomic DNA. For hPGCLC and oogonia generation, polyclonal pools of hiPSCs were utilized. Average copy number was 8-10. The same procedure was performed for generating combinatorial cell lines, in which the 100 fmol of cDNA vector was divided equally between each TF for a pooled nucleofection. For copy number assessment the following was performed: 1) RT-qPCR using SYBR Green master mix was performed after gDNA extraction using the DNAeasy kit. 2) 10 ng of input gDNA was used per reaction based on the standard curve, with an anneal temperature of 60 degrees. 3) To calculate copy number, the 2^ΔCq+1^ method was used, with RNAseP as a reference. 4) The resultant value was multiplied by two to account for the two autosomal copies of *RPP30*.

### Generation of NANOS3+ and DDX4+ cells in 2D Monolayer

For generation of NANOS3+ and DDX4+ cells in monolayer, an identical induction format and media composition was utilized. hiPSCs containing integrated TF expression cassettes were cultured in mTeSR1 medium on Matrigel coated plates. For induction in monolayer, hiPSCs were dissociated to single cells using Accutase and seeded onto Matrigel coated plates at a density of 2,500–3,000 cells per cm^2^ in mTeSR1 + 10μM Y-27632 and 1μg/ml doxycycline for 6 hours. Media was then removed and washed with DMEM/F12 and replaced with Media 1 (see components list below). After 12-18 hours of induction, Media 1 was removed and washed with DMEM/F12 and replaced by Media 2. After 24 hours, Media 2 was removed and replaced by Media 3. After 24 hours, Media 3 was replaced with Media 4. Reporter positive cells could then be harvested for use after 24 hours in Media 4. Putative hPGCLCs could be isolated via the NANOS3 reporter expression, CD38 cell surface expression, combinations of both or EpCAM/ITGA6 dual positive cell surface markers. Induced oogonia-like cells (iOLCs) could be isolated via a DDX4 reporter. hPGCLCs and iOLCs could additionally be generated via embryoid body formation through methods established in Irie et al. 2015, Sasaki et al. 2015, Yamashiro et al. 2018, Kobayashi et al. 2022, and Murase et al. 2021.

Media formulations were as follows: **Basal media (aRB27):** Advanced RPMI, 1X B27 minus Vitamin A, 1X Glutamax, 1X Non-Essential Amino Acids, 10 µM Y-27632, 1X Primocin or Pen-Strep, 1 µg/ml Doxycycline. **Media #1 (Epiblast-induction media):** aRB27 Basal Media, 3 µM CHIR99021, 100 ng/ml Activin A, 0.1 μM PD173074. **Media #2**: aRB27 Basal Media, 1 μM XAV939, 40 ng/ml hBMP4. **Media #3:** aRB27 Basal Media, 1 μM XAV939, 100 ng/ml SCF, 50 ng/ml EGF. **Media #4**: aRB27 Basal Media, 1 μM XAV939, 40 ng/ml hBMP4, 100 ng/ml SCF, 50 ng/ml EGF.

### Fluorescent Activated Cell Sorting of hPGCLCs and iOLCs

hPGCLCs and iOLCs were analyzed using flow cytometry on the BD LSRFortessa or Cytoflex LX machine. Cell events were selected on FSC-A versus SSC-A, singlets chosen via FSC-W versus FSC-H dispersion, and live cells selected based on DAPI negative gating on the Indo-Violet-A channel (Figure S9). Negative gates were set using hiPSC controls or unstained cell controls. For any multi-staining flow cytometry, single stain controls and beads were utilized for compensation. For cell sorting for post-isolation growth, cells were captured using a Sony SH800S cell sorter. All flow cytometry analysis was conducted using the FlowJo software system (v10.8.1). NANOS3 and DDX4 were captured via fluorescent reporters. For cell surface markers, CD38 PerCP-Cy5.5 Mouse IgG (Biolegend 303522), EpCAM-APC-Cy7 Mouse IgG (Biolegend 324245), and Integrin-alpha-PE Rat IgG (Biolegend 313607) were utilized. A 1:60 dilution of each antibody in a dPBS + 3% FBS FACS buffer was utilized. Staining was performed for one hour at 4 °C after harvesting with Accutase, and cells were sorted without fixation with additional DAPI staining.

### Barcode enrichment screening for additional factors

Pooled PiggyBac plasmid libraries (Figure 4A/B) were integrated into 1 million hiPSCs per replicate, using a Lonza nucleofector with 100 µL of P3 buffer. 250 fmol transposon pool and 2.5 µg transposase plasmid were used for each nucleofection. Puromycin selection was performed between days 2-7 post-nucleofection. Subsequently, DDX4+ iOLCs were induced using the protocol described above. In followup rounds of screening, cells were treated with 5 µM GSK-3484862, and some replicates were additionally treated with the retinoic acid receptor agonist AM580 (1 µM). DNA was extracted from sorted DDX4+ cells using the Qiagen DNEasy Micro kit, and from unsorted and pre-induction cells using the Qiagen DNEasy kit. Barcode amplification and sequencing was performed as previously described.^51^ For fold-change calculations, barcode abundances in the DDX4+ cells were separately compared to unsorted cells and to pre-induction cells.

### Improved protocol for iOLC induction

Human or rhesus iPSCs were dissociated using Accutase and seeded onto Matrigel coated plates at a density of 50,000/cm2 in cold mTESR1 supplemented with 2% Cultrex (R&D Systems), 10 µM Y-27632, 5 µM GSK-3484862, and 1 µg/mL doxycycline. After 24 hours, cells were washed with RPMI and the medium was changed to aRB27 with 25 ng/mL hBMP4, 2% Cultrex, 5 µM GSK-3484862, and 1 µg/mL doxycycline. After 24 hours, the cells were fed with fresh medium. After 24 hours, the medium was changed to aRB27 with 50 ng/mL SCF, 50 ng/ml EGF, 10 ng/mL hLIF, 5 µM GSK-3484862, and 1 µg/mL doxycycline. After 24 hours, the cells were fed with fresh medium. After 24 hours, cells were harvested with Accutase for downstream analysis.

### Transient naïve resetting of hiPSCs

Naïve resetting was performed as previously described (MacCarthy et al., 2024). Briefly, 3 μg of episomal pCXLE-Sox2^A61V^-2A-Klf4-2A-Myc (Addgene #210017) and 3 μg of pCXWB-EBNA1 (Addgene #37624) were co-nucleofected into 1×10^6^ primed hiPSCs using the Lonza 4D-nucleofector (“Primary Cell P3” solution, “CM-113” program) and plated on a feeder layer of irradiated CF1 MEFs (Thermofisher, 4×10^6^ MEFs per 6 well plate) at 8×g10^4^ nucleofected cells per well, in StemFlex media supplemented with 10 µM Y-27632. The next day, the medium was changed to PXL medium. The cells were fed daily. At day 6, the cells were harvested using Accutase, and used for iOLC induction (without sorting).

Composition of PXL medium: 1:1 mix of Neurobasal medium (Gibco, 21103049) and Advanced DMEM/F12 (Gibco, 11320082) supplemented with 1X N2 (Gibco, 17502048), 1X B27 minus vitamin A (Gibco, 12587010), 1X sodium pyruvate (Gibco, 11360070), 1X non-essential amino acids (Gibco, 11140050), 1X GlutaMAX (Gibco, 35050061), 1X Penicillin-Streptomycin (Gibco, 15070063), 0.1 mM b-mercaptoethanol (Gibco, 31350010, 1ml in 500), 50 μg/ml L-ascorbic acid (Sigma, A8960), 0.2% Geltrex (Gibco, A1413301), 1 μM PD0325901, 2 μM XAV939 (Sigma, X3004), and 20 ng/ml hLIF (Peprotech, 300-05).

### Enzymatic methylation sequencing

Genomic DNA was extracted from hiPSCs, hPGCLCs, and sorted DDX4+ iOLCs using the Qiagen DNeasy Micro kit. Libraries were prepared from 200 ng DNA using the NEB Enzymatic Methyl-Seq kit (NEB #E7120) following the manufacturer’s instructions. Briefly, control DNA (methylated pUC19 and unmethylated λ phage) was spiked into each sample. Next, samples were sheared to ∼300 bp using a Covaris S2 sonicator. Enzymatic methyl conversion was performed, and libraries were amplified using unique dual index primers. 2 x 150 bp paired end sequencing was performed on an Illumina NovaSeq.

### Methylation analysis

Reads were aligned using bismark (v 0.24.0)^56^ to the hg38 reference genome, with additional pUC19 and λ phage sequences for analysis of the control DNA. For analysis of methylation at functional loci, CpGtools CpG_aggregation.py ^57^ was used to aggregate methylation for features of interest, including promoters (defined as 900bp upstream and 400bp downstream of all transcriptional start sites), imprint control regions, CpG islands, and transposable elements. With the exception of imprint control regions (obtained from Jima *et al.* 2022),^43^ genomic coordinates of all features were obtained from the UCSC hg38 TableBrowser database (https://genome.ucsc.edu/cgi-bin/hgTables).^58^

Identification of differentially methylated regions was performed using the R package DSS (version 2.50.1) with default settings.^59^ As DSS only calculates a false discovery threshold for individual loci, not regions, we performed a permutation analysis to identify a suitable areaStat threshold, which we set at 600. Annotation of differentially methylated regions was performed using `bedtools intersect` (version 2.31.0)^60^ with the feature lists described above. For making plots, features were colored in reverse order of abundance (promoters > ICRs > CpG islands > TEs).

### Transcriptomic characterization of TF induced cell lines

Cells were induced according to the above protocols and isolated for bulk RNA-sequencing. Library preparation was performed using the NEB Next ultra-low input RNA-sequencing library preparation kit with PolyA capture module for samples containing less than 10,000 cells. For samples with greater than 10,000 cells the NEBNext Ultra II RNA-sequencing library preparation kit with PolyA mRNA capture module was utilized. Sequencing was performed on Illumina Next-Seq 500 and NovaSeq platforms with 2 x 100bp or 2 x 150bp paired end reads, respectively. Only samples with RNA quality RIN scores of greater than 8 were utilized for analysis.

For DLX5, HHEX, and FIGLA hPGCLC bulk RNA-sequencing, TFs were induced for four days and cells were sorted for NANOS3+ expression and utilized for bulk RNA-Seq. No-TF control cells were also utilized and sorted via NANOS3. For long term culture (LTC) hPGCLCs, data was downloaded from GEO and realigned using our pipeline. For D3 DDX4+ bulk RNA-Seq, TFs were induced for four days and cells were sorted for DDX4+ expression and utilized for bulk RNA-Seq. For reference samples of the ovarian atlas, data was downloaded from GEO and realigned using our pipeline. The reference samples used are indicated in Supplementary Table 3.

### RNA-Seq Analysis

Reads from our samples as well as previously published datasets were aligned to the human reference genome (GRCh38) using STAR (version 2.7.6),^61^ to construct count matrices aligning sequencing reads to the known set of human genes. For samples with 2 x 150 bp reads, the STAR options `--outFilterScoreMinOverLread 0.33--outFilterMatchNminOverLread 0.33` were used in order to keep the minimum match length equivalent between the different samples. After alignment, reads were normalized using DESeq2’s median of ratios.^52^ A heatmap was generated in R (version 4.3.2) using ComplexHeatmap. Differential expression analysis was performed using DESeq2 (version 1.42.0). For comparisons between different studies, pluripotent stem cell samples (our hiPSCs, Yamashiro *et al* hiPSCs, and Tang et al hESCs) were labeled as “hPSCs”, and each study was added as a covariate to the DESeq2 model. For each comparison, lists of significantly (p_adj_ < 0.05) upregulated (log2fc > 1) and downregulated (log2fc < −1) genes were used as input for GO term enrichment analysis with the PantherDB API (version 18.0).^62^

### Floating aggregate hPGCLC induction and harvesting

hiPSCs were differentiated to hPGCLCs according to the method of Sasaki et al. 2015.^19^ Briefly, iMeLCs were induced by plating hiPSCs on matrigel coated plates in GK15 medium (GMEM with 15% KSR, 0.1 mM NEAA, 2 mM L-glutamine, 1 mM sodium pyruvate, and 0.1 mM 2-mercaptoethanol) containing Activin A, CHIR, and ROCK inhibitor (Y-27632). The hPGCLCs were induced by plating iMeLCs into a well of a low-cell-binding V-bottom 96-well plate in GK15 supplemented with LIF, BMP4, SCF, EGF, and ROCK inhibitor for 4 days.

For PGCLC harvesting and staining for FACS, for each sample, nine aggregates were combined in a 1.5 mL tube. The aggregates were washed with PBS, resuspended in trypsin solution (45 µL), and incubated in a shaking heat block (37 °C, 800 rpm). After 8 minutes, 940 µL FACS buffer (3% FBS in PBS) was added, and the aggregates were dissociated by vigorous pipetting. The suspension was passed through a 70 µm strainer and spun down (300 *g*, 3 min). The supernatant was removed, and the pellet was resuspended in antibody solution (40 µL). The suspension was kept on ice in the dark for 30 minutes. Then, 940 µL FACS buffer was added. The cells were spun down again and resuspended in 200 µL FACS buffer with 100 ng/mL DAPI, and analyzed on a BD LSR Fortessa flow cytometer as described above.

### iOLC maintenance culture

Feeder-free maintenance of iOLCs post-isolation was accomplished on Matrigel coated plates with growth in S-CM media, established by Kobayashi et al. 2022, which is an STO-feeder cell conditioned media supplemented with SCF. The iOLC expansion protocol is identical to the expansion protocol for long term culture of hPGCLCs and shows maintenance of DDX4+ expression over 28 days. The hPGCLC basal medium contained 13% (v/v) KSR, 1x NEAA, 1 mM sodium pyruvate, and 1x penicillin-streptomycin in Glasgow’s MEM with 2 mM glutamine (ThermoFisher, 11710035). STO-CM was prepared by maintaining 5.0e6 mitomycin C treated STO cells in 12 mL of hPGCLC basal medium for 24 h, removing cells by centrifugation, and storing frozen at −20°C until use. The complete hPGCLC maintenance medium (S-CM for SCF-supplemented CM) was prepared by adding 0.1 mM b-mercaptoethanol, 50 mg/mL L-ascorbic acid, and 100 ng/mL recombinant human SCF to the CM. FACS-enriched DDX4+ iOLCs (100-5000 cells) were inoculated onto Matrigel coated plates in a well of six-well plates (Corning, 3506) with SCF-supplemented hPGCLC basal medium containing 10 µM Y27632 + 1 μg/ml doxycycline. Medium was changed every other day without Y27632. During the iOLC expansion, cells were dissociated using Accutase and passaged onto fresh matrigel every 5–14 days, when cells reached 50% confluency.

### Immunofluorescence imaging

Cells cultured on Matrigel-coated ibidi 8-well plates (iOLCs) (ibidi, cat 80806) were washed once with 200 μL dPBS and fixed by treatment with 200 μL of 4% paraformaldehyde in dPBS for 10 minutes at room temperature. The dPBS wash was repeated twice, and the cells were permeabilized by treatment with 200 μL 0.25% Triton X-100 in dPBS for 10 minutes at room temperature. The cells were washed with 200 μL PBST (0.1% Triton X-100 in dPBS), and blocked with 100 μL blocking buffer (1% bovine serum albumin and 5% normal donkey serum [Jackson ImmunoResearch, cat 017-000-121, lot 152961] in PBST) for 30 minutes at room temperature. The blocking buffer was removed and replaced with a solution of primary antibodies in blocking buffer, and the cells were incubated overnight at 4°C. The antibody solution was removed and the cells were washed three times with 200 μL PBST. The cells were incubated with 100 μL secondary antibody solution in blocking buffer for 1 hour at room temperature in the dark. The secondary antibody was removed and replaced with 200 μL of DAPI solution (1 ng/mL in dPBS). After 10 minutes the DAPI solution was removed and the cells were washed twice with 200 μL dPBS, and stored in dPBS at 4°C in the dark until imaging (typically a few hours).

hPGCLC floating aggregates were washed with PBS and fixed with 1% PFA overnight at 4°C. After another PBS wash, aggregates were detached from the Transwell. In preparation for cryosectioning, and transferred to 10% sucrose in PBS. After 24 hr at 4°C, the 10% sucrose solution was removed and replaced with 20% sucrose in PBS. After an additional 24 hr at 4°C, the ovaroids were embedded in OCT compound and stored at −80°C until sectioning.

The aggregates were sectioned to 10 μm using a Leica CM3050S cryostat. Sections were transferred to Superfrost Plus slides, which were washed with PBS to remove OCT compound. The slides were washed with PBST (0.1% Triton X-100 in PBS) and sections were circled with a Pap pen. Slides were blocked for 30 min at room temp. with blocking buffer (1% bovine serum albumin and 5% normal donkey serum in PBST). The blocking buffer was removed and replaced with a solution of primary antibodies in blocking buffer, and the slides were incubated overnight at 4°C. The antibody solution was removed and the slides were washed with PBST for 3 times for 5 minutes. The slides were incubated with secondary antibody and DAPI solution in blocking buffer for 1 hr. at room temp. in the dark, followed by two 5 minute washes with PBST and one wash with PBS. After staining, samples were mounted in Prolong Gold medium and covered with coverslips. Imaging was performed on a Leica SP5 confocal microscope.

Imaging was performed on a Leica SP5 confocal microscope. Antibodies used are listed as follows: **Primary Antibodies:** OCT4-Mouse IgG (AB398736) - 1:250 Dilution, DDX4-Mouse IgG (AB27591) - 1:500 Dilution, DAZL-Rabbit IgG (AB215718) - 1:167 Dilution, SOX17-Goat IgG (AB355060) - 1:500 Dilution, ITGA6-Rat IgG (AB493634) - 1:167 Dilution (1:60 for flow cytometry). **Secondary Antibodies:** Mouse IgG-AF647 Donkey (AB162542) - 1:250 Dilution, Rabbit IgG-AF488 Donkey (AB2534104) - 1:500 Dilution, Goat IgG-AF568 Donkey (AB2534104) - 1:500 Dilution.

### Quantification and Statistical Analysis

All statistical analysis and specific quantification can be found in the figure legends and methods sections. All graphing was performed using GraphPad unless otherwise noted. For Figure 2B, n=3 replicates were independently seeded for induction from a single hiPSC cell line. Figure 2C, n=6 replicates were independently seeded for induction from three independent hiPSC cell lines, two per line. Results from each cell line were averaged for the duplicates and graphed as biological replicates. For Figure S4, n=3 replicates were independently seeded for induction from a single hiPSC cell line. For Figure S5, n=3 replicates were generated from a single hiPSC cell line by pooling n=9 independent floating aggregates.

## Supplementary Figures

Figure S1. Expression of prioritized transcription factors in the human fetal gonad atlas

Figure S2. Development of dual fluorescent reporter hiPSC line for TF screening

Figure S3. Development of doxycycline inducible TF screening cell lines

Figure S4. Single TF overexpression screening for NANOS3+ cell yield

Figure S5. *DLX5*, *HHEX,* and *FIGLA* overexpression significantly improves hPGCLC yield

Figure S6. TF-derived hPGCLCs display canonical protein expression in floating aggregate differentiation

Figure S7. Additional DNA methylation analysis of iOLCs, hPGCs, and xrOvary cells.

Figure S8. Principal component analysis of gene expression. Figure S9. Representative flow cytometry analysis

Supplementary Table 1: PageRank Ranking of all TFs

Supplementary Table 2: TFs utilized for screening

Supplementary Table 3: Normalized gene counts of all RNA-seq samples

Supplementary Table 4: DESeq2 log fold changes and adjusted p-values

Supplementary Table 5: Significantly enriched GO terms

Supplementary File 2: Monolayer germ cell screening reagents and methods

## Author Contributions

C.C.K. and P.C. conceived, designed, and directed the study. P.C. developed and implemented GRN inference and centrality analysis algorithms. C.C.K. performed TF screening, cell line creation, differentiation protocol development, flow cytometry, cell engineering, iOLC expansion culture methods, and sequencing library preparation. M.P.S. generated reporter cell lines, performed live cell imaging, aided in flow cytometry, aided in TF screening, and conceived and developed the improved iOLC induction protocol. Se.V. performed transient naïve resetting. C.M. aided in barcode enrichment analysis. P.R.F. performed confocal imaging, cell line expansion, aided in cell culture and library preparation. B.W. assisted with library preparation and analysis. P.C., V.S.K., M.P.S., S.G., So.V., and T.C. conducted RNA-seq analysis. M.P.S. and U.W. conducted methyl-seq analysis. E.D. and J.A. assisted in vector cloning, TF screening, cell culture, and confocal imaging. M.K. performed fetal gonad TF expression analysis. M.P.S., C.C.K., and P.C. wrote the manuscript. P.C. supervised the study, with assistance from R.E.K., T.S., and G.M.C.

## Data and Materials Availability

All data needed to evaluate the conclusions in the paper are present in the paper and supplementary tables and figures. Data analysis code can be found at: https://github.com/programmablebio/egg. Raw and processed sequencing data will be deposited to GEO upon publication.

## Acknowledgements

This work was funded by the Synthetic Biology Platform at the Wyss Institute for Biologically Inspired Engineering and institutional startup funds at Duke University. We thank Songlei Liu, Sabrina Koseki, Kalyan Palepu, Garyk Brixi, Suhaas Bhat, Emma Tysinger, and Teodora Stan for technical assistance in the research. This work was additionally funded by the Gameto Sponsored Research Agreement at Harvard University and Colossal Sponsored Research Agreement at Harvard University. M.P.S. was supported by the National Science Foundation Graduate Research Fellowship and NICHD F31 Fellowship (F31HD108898-01A1).

