## Supplementary Information for "Rapid Human Oogonia-like Cell Specification via Combinatorial Transcription Factor-Directed Differentiation"

**Supplementary Figures**

1. Expression of prioritized transcription factors in the human fetal gonad atlas
2. Development of dual fluorescent reporter hiPSC line for TF screening
3. Development of doxycycline inducible TF screening cell lines
4. Single TF overexpression screening for NANOS3+ cell yield
5. *DLX5*, *HHEX,* and *FIGLA* overexpression significantly improves hPGCLC yield
6. TF-derived hPGCLCs display canonical protein expression in floating aggregate differentiation
7. D3-derived DDX4+ cells transcriptomically resemble fetal oogonia
8. Representative flow cytometry analysis

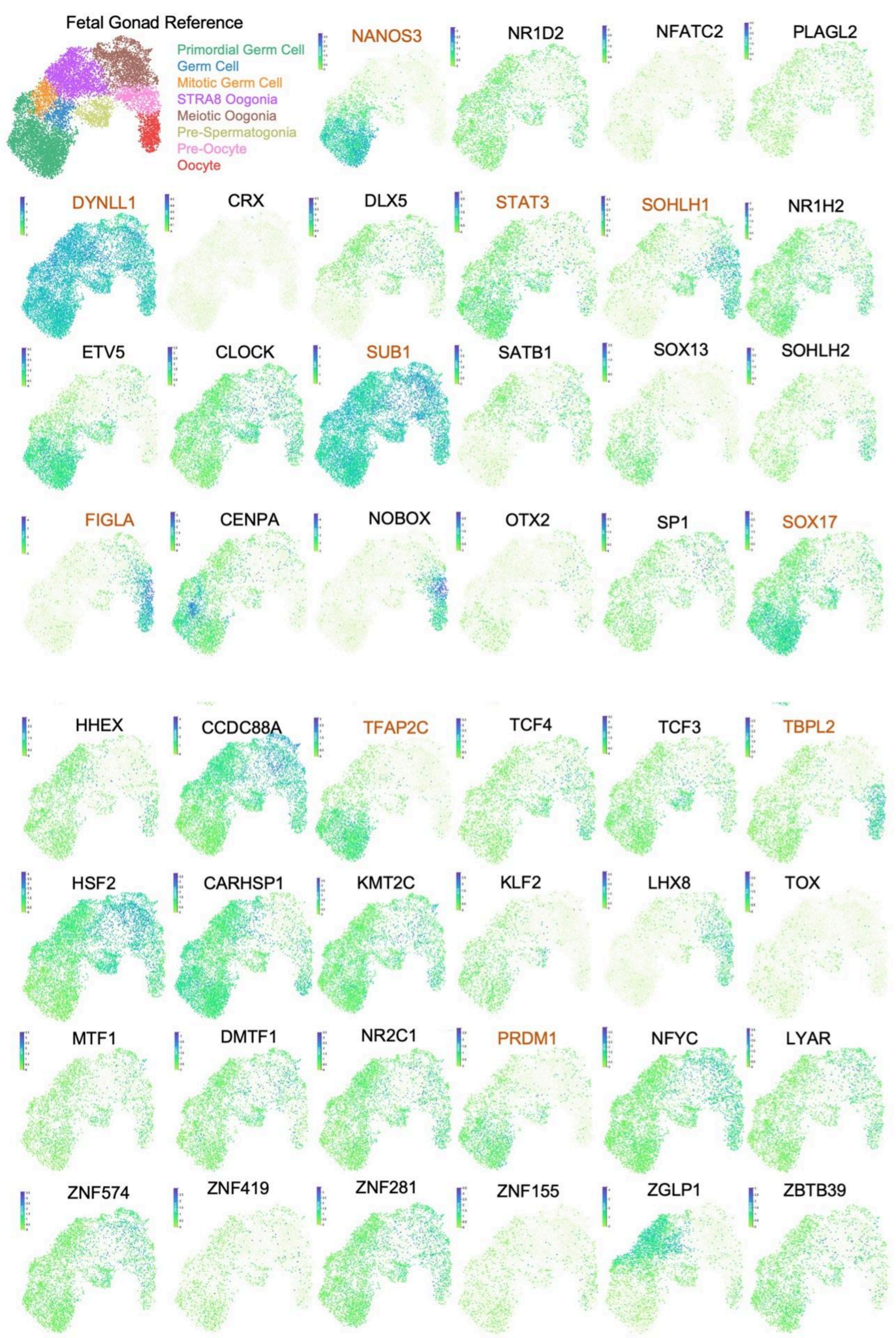


**Supplementary Figure 1.** Expression of prioritized transcription factors in the human fetal gonad atlas**, Related to Figure 1.** The expression of all 46 TFs was visualized using the reproductive cell atlas, utilizing only germline cells obtained from the fetal atlas of Garcia-Alonso et al. 2022. Expression level for each TF is mapped onto the UMAP, with the cell type annotation shown in the reference map. TFs included as controls in the study are highlighted orange.

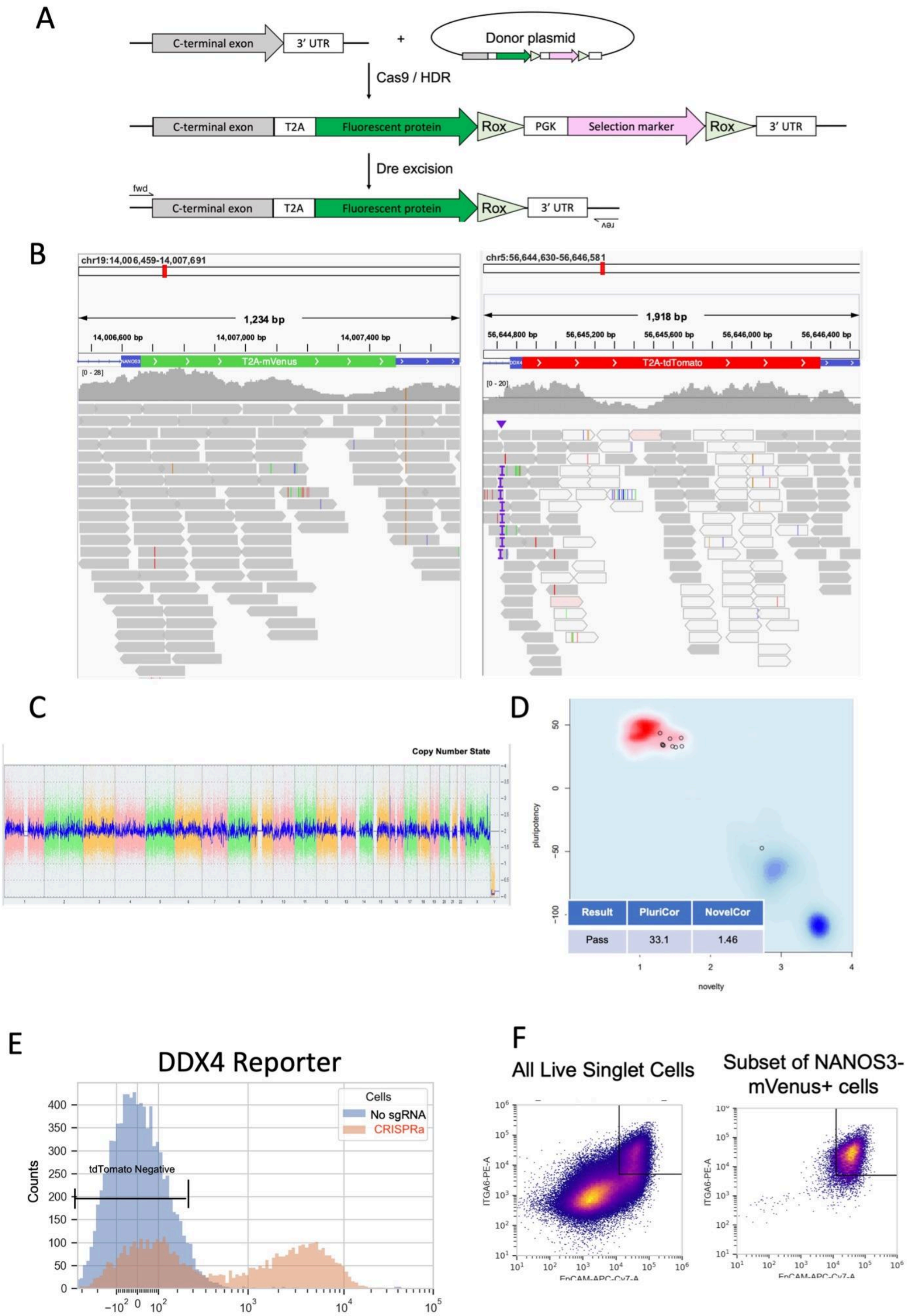


**Supplementary Figure 2.** Development of dual fluorescent reporter hiPSC line for TF screening**, Related to Figure 2.** A) Schematic representation of cell line generation methodology. B) Genotyping WGS of the reporter line utilized in study, with specific display of the inserted reporter construct regions. C) The whole genome view displays all somatic and sex chromosomes in one frame with high level copy number. The smooth signal plot (right y-axis) is the smoothing of the log2 ratios which depict the signal intensities of probes on the microarray. A value of 2 represents a normal copy number state (CN = 2). A value of 3 represents chromosomal gain (CN = 3). A value of 1 represents a chromosomal loss (CN = 1). The pink, green and yellow colors indicate the raw signal for each individual chromosome probe, while the blue signal represents the normalized probe signal which is used to identify copy number and aberrations D) The pluripotency plot window provides a visual representation of the tested samples in the analysis. The pluripotency and novelty x/y scatter plot combines the pluripotency score on the y-axis with the novelty score on the x-axis. The red and blue background hint to the empirical distribution of the pluripotent (red) and non-pluripotent (blue) samples in the reference data set. The samples were analyzed using an algorithm that integrates gene expression data to authenticate pluripotency status. Samples are screened against samples in the stem cell database and given a pluripotency score (PluriCor) and novelty score (NovelCor), which are shown in the table. Pass shows a clear pluripotency signature. Fail means the samples are not pluripotent. A non-iPSC sample was used in this experiment to serve as a negative control for non-pluripotency. E) Flow cytometry analysis after induction of the DDX4-reporter locus through CRISPR-activation, blue represents a no sgRNA control and red represents the CRISPRa- sgDDX4 condition, the bracket highlights the cells considered reporter negative. F) Flow cytometry analysis of hPGCLC formation using the method used for screening (Figure 2A) for the D4TN3V reporter line. Live, single cells are visualized for expression of the cell surface markers EpCAM and ITGA6 (left panel). Additionally in the subset of NANOS3-mVenus+ cells, expression of EpCAM/ITGA6 is visualized (right panel).

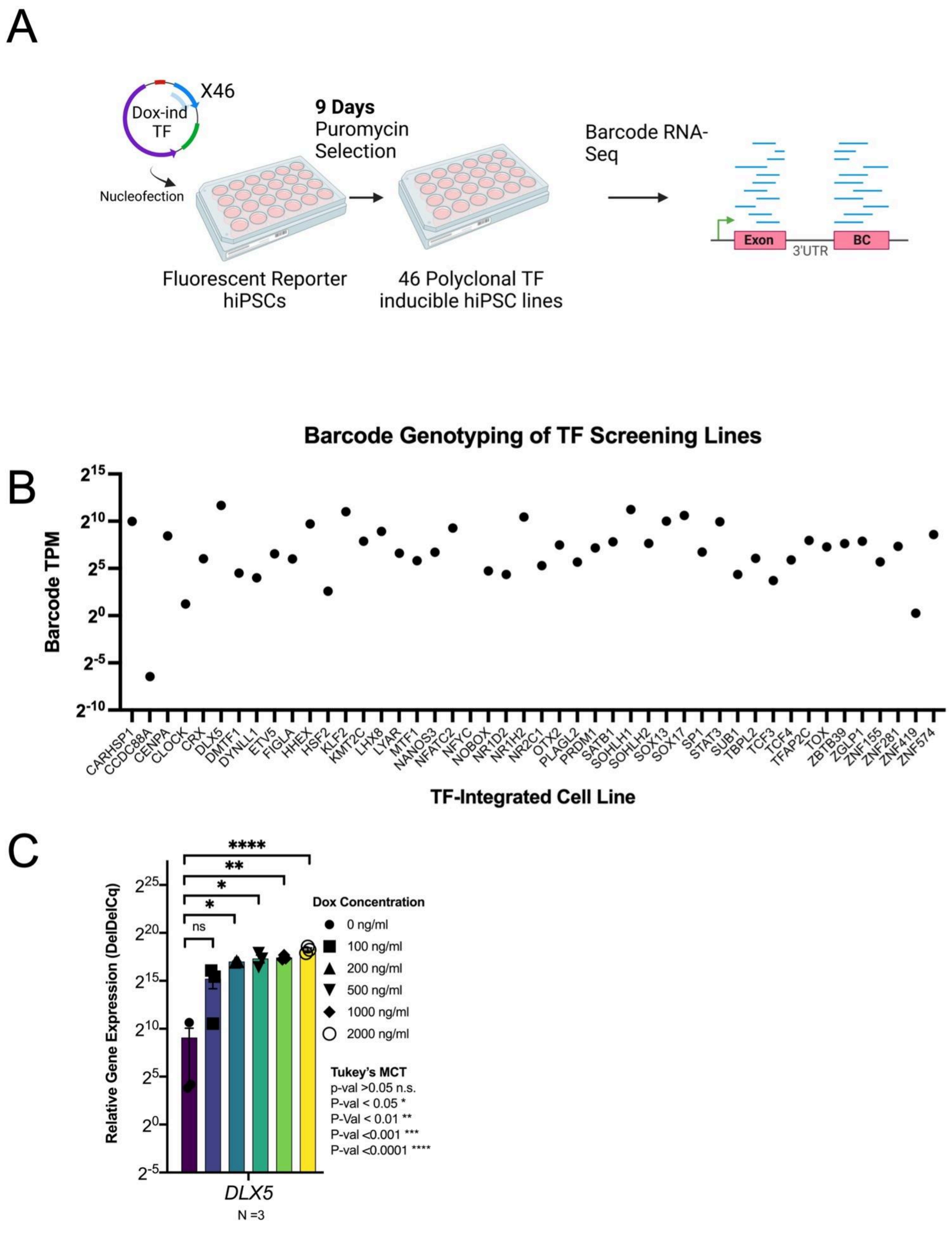


**Supplementary Figure 3.** Development of doxycycline inducible TF screening cell lines**, Related to Figure 2.** Schematic representation of 46 cell line generation B) Genotyping via barcode capture through RNA-seq. Transcriptomes were captured and analyzed via Kallisto for barcode identification. Barcode transcripts per million (TPM) from Kallisto analysis of 46 cell lines. The Barcode TPM for each intended TF is visualized for each line. C) RT-qPCR of the *DLX5* gene under doxycycline serial dilution in three independent replicates plotted relative to GAPDH using the deldelCq method. Statistical analysis is performed using ANOVA with multiple comparison testing.

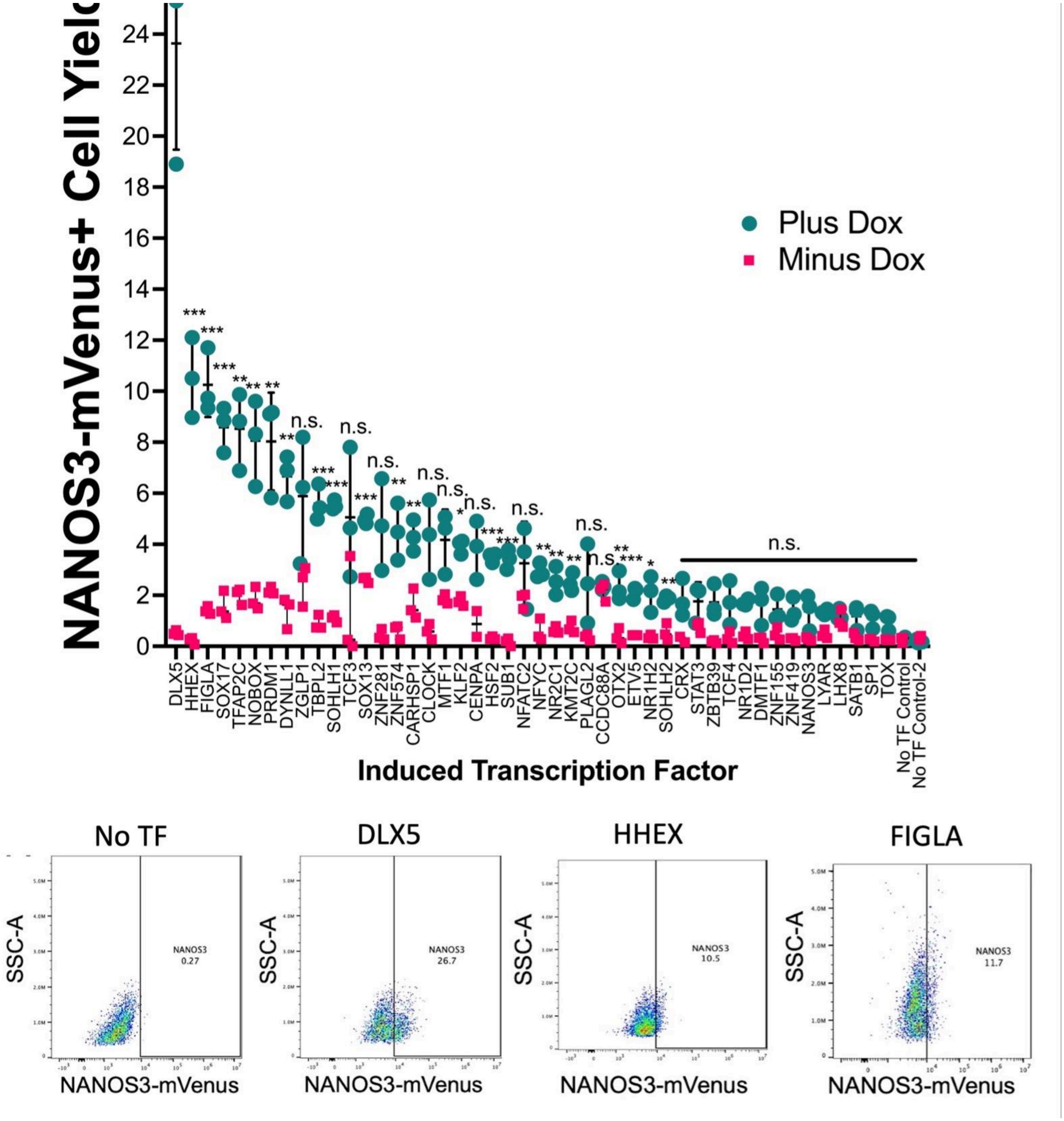


**Supplementary Figure 4.** Single TF overexpression screening for NANOS3+ cell yield**, Related to Figure 2.** Results for NANOS3-T2A-mVenus flow cytometry for triplicate induction conditions in the plus (blue) and minus (pink) doxycycline induction condition for each TF. Data is plotted as a percent of NANOS3-T2A-mVenus+ cells on the y-axis for each TF. Individual dots represent individual wells of the induction condition, seeded separately from a single hiPSC line. Horizontal black line represents the mean of induction replicates. Statistical significance was determined by multiple T-test comparison between the plus and minus dox condition for each TF, with a p-value <0.05 considered as significant. The FDR correction was utilized for multiple hypothesis testing. *** p<0.001, ** P<-.01, * p<0.05. Representative flow cytometry plot are shown for the No TF control condition and the three highest yielding conditions: DLX5, HHEX, and FIGLA.

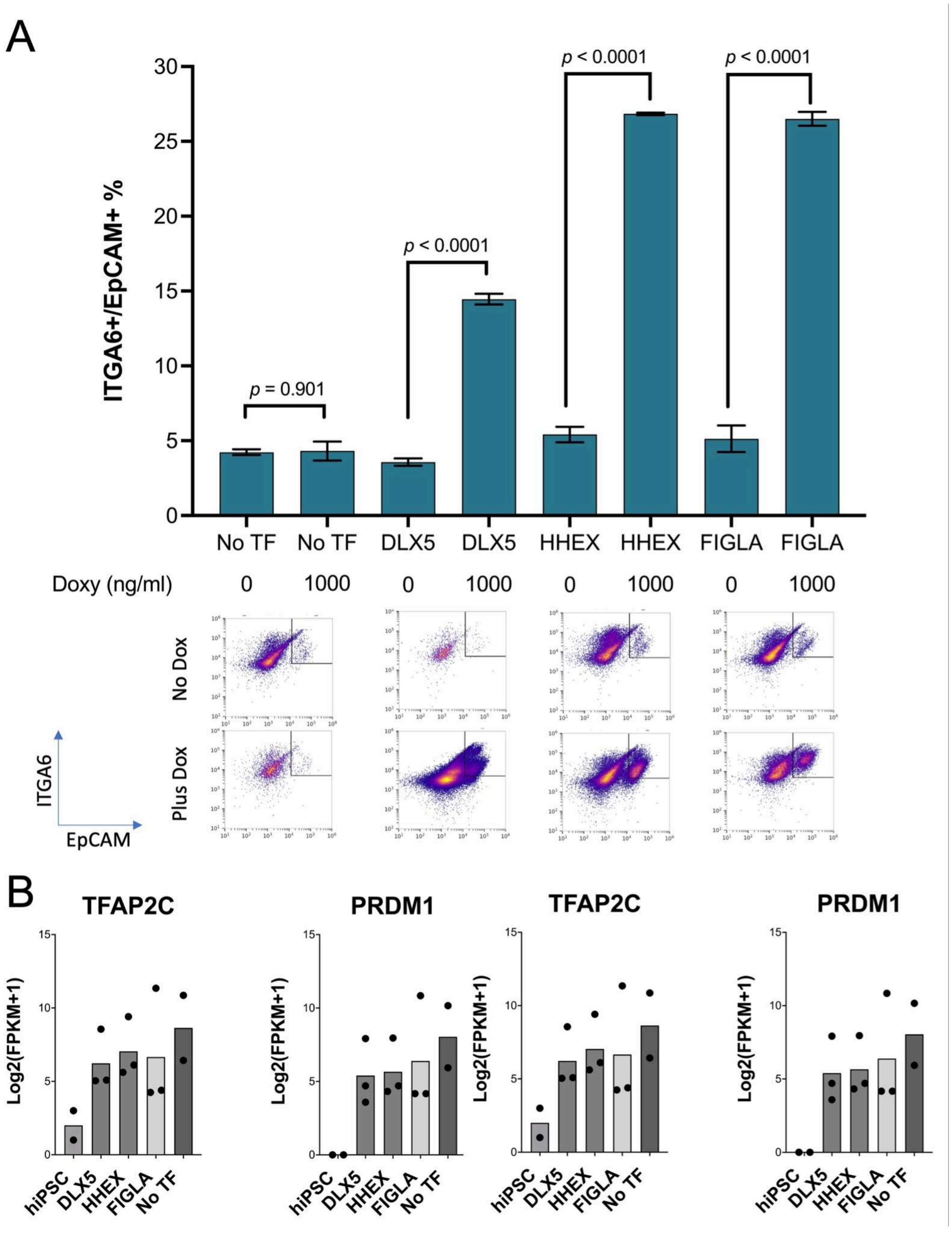


**Supplementary Figure 5.** *DLX5*, *HHEX,* and *FIGLA* overexpression significantly improves hPGCLC yield**, Related to Figure 3.** A) Generation of hPGCLCs in N = 3 replicates of n=9 pooled embryoid bodies per condition. hPGCLCs were induced according to the protocol from Sasaki et al. 2015 and analyzed via flow cytometry for the markers EpCAM and ITGA6. hPGCLCs were induced in the presence and absence of doxycycline in lines harboring inducible vectors for *DLX5*, *HHEX*, and *FIGLA* and a no TF control*.* Statistical analysis was performed via unpaired two-tailed *t*-test between doxycycline and no doxycycline conditions for each condition. Representative flow cytometry graphs are shown for EpCAM/ITGA6 on live singlets for the no dox and plus dox conditions B) Expression of known hPGCLC marker genes (*SOX17*, *TFAP2C*, *PRDM1*) and hiPSC marker gene (*SOX2*), in NANOS3+ sorted cells isolated from monolayer induction in the no TF control, *DLX5*, *HHEX*, and *FIGLA* overexpression conditions. RNA abundance was measured via RNA-seq for N=2 or 3 independent replicates and plotted via Log2 normalization of FPKM +1 values against an hiPSC control.

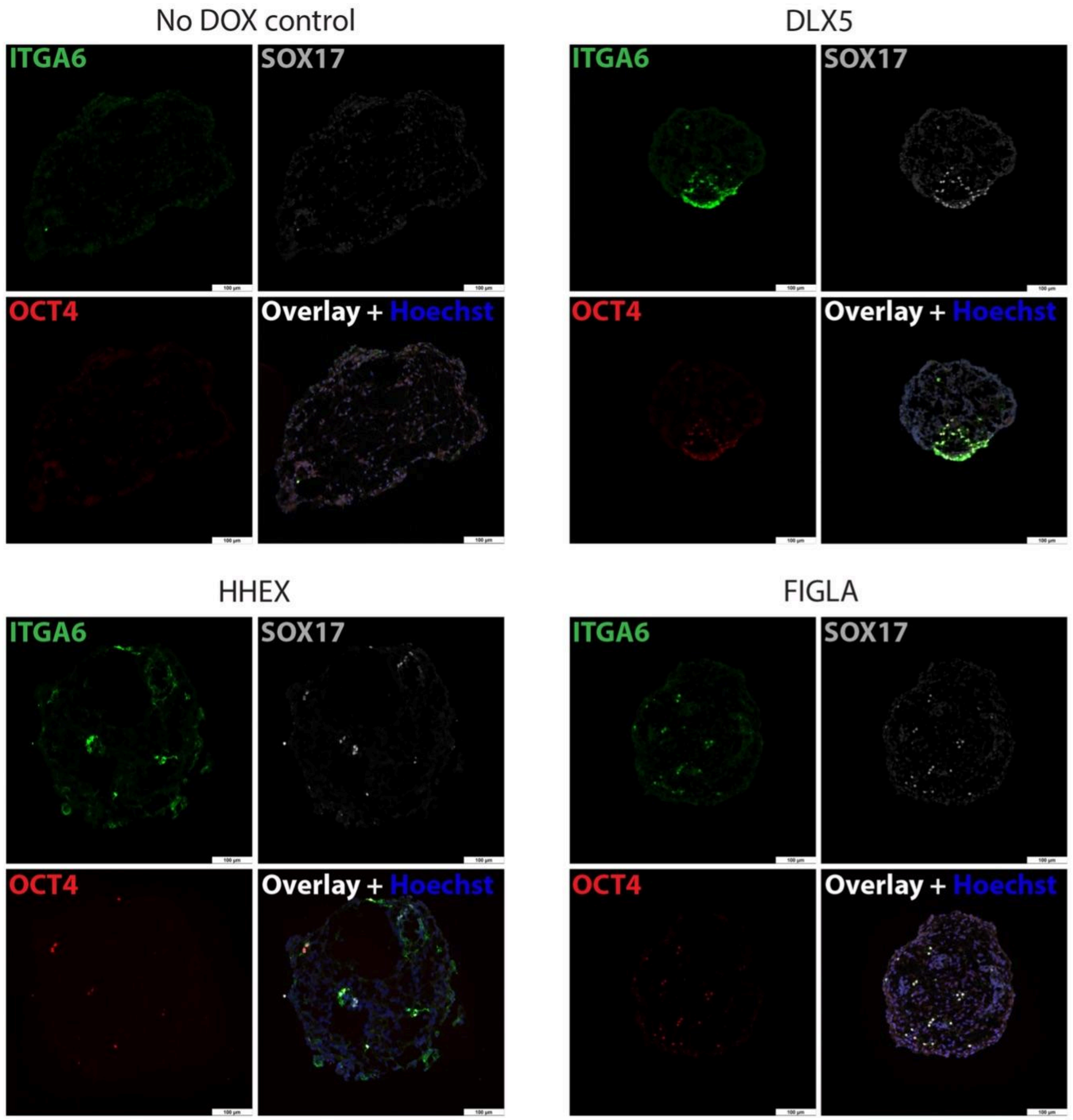


**Supplementary Figure 6.** TF-derived hPGCLCs display canonical protein expression in floating aggregate differentiation**, Related to Figure 3.** hPGCLCs were induced according to the protocol from Sasaki et al. 2015 and analyzed via immunofluorescence after fixation and cryosectioning. hPGCLC markers ITGA6 (green), SOX17 (red), and OCT4 (grey) are visualized alongside Hoechst staining (blue). A no doxycycline control and embryoid bodies from *DLX5*, *HHEX*, and *FIGLA* inductions are shown at day 4 of differentiation. Scale bar is 100μm.

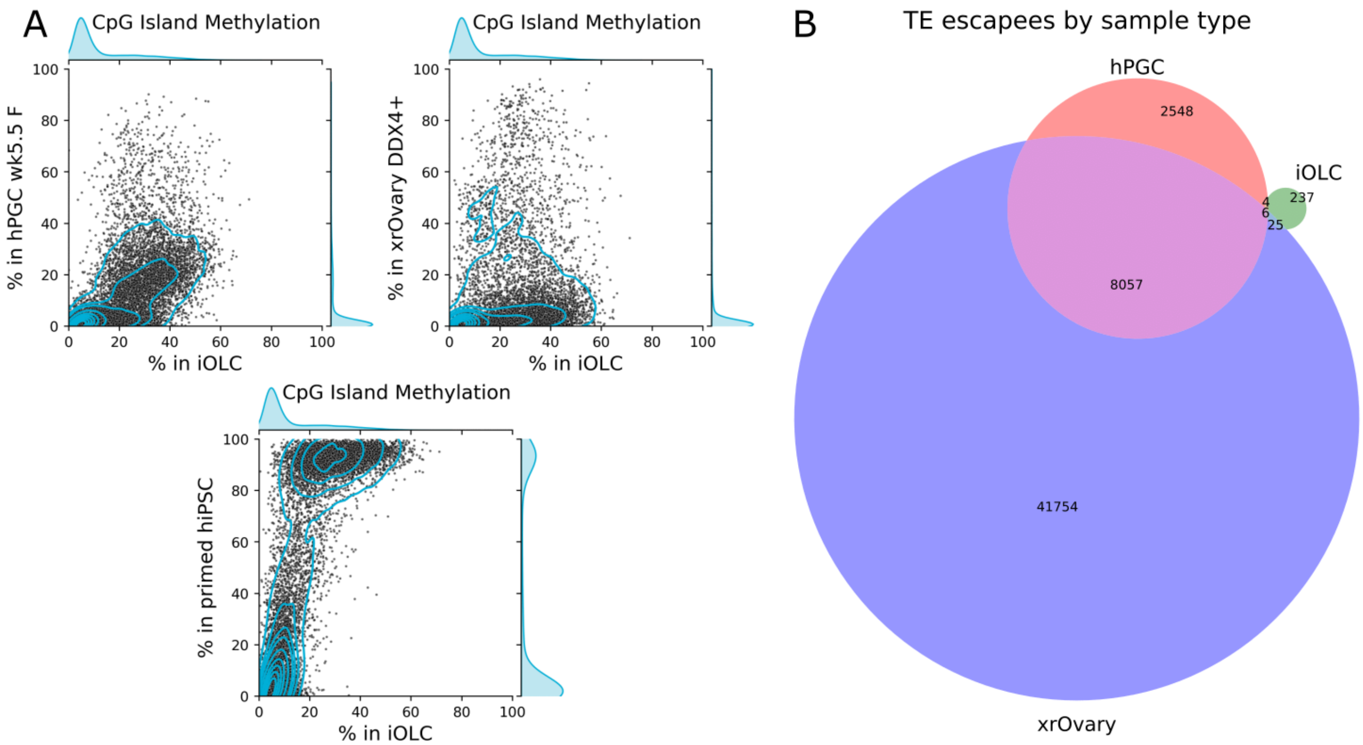


**Supplementary Figure 7.** Additional DNA methylation analysis of iOLCs, hPGCs, and xrOvary cells (related to Figure 5). (A) CpG island methylation in primed hiPSCs, iOLCs (induced using D5 TFs and the improved protocol), week 5.5 female hPGCs, and xrOvary day 120 DDX4+ cells. (B) Overlap of TEs escaping demethylation (defined as >80% average methylation) in iOLCs, week 5.5 female hPGCs, and xrOvary day 120 DDX4+ cells.

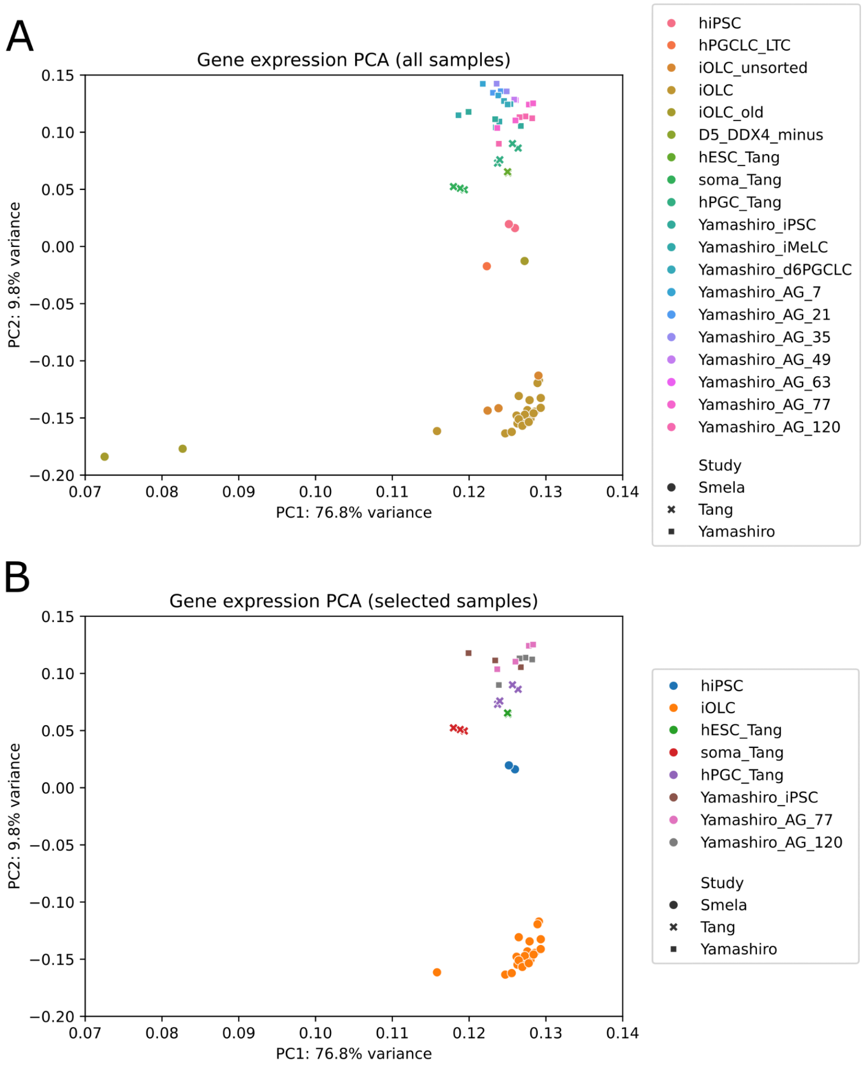


**Supplementary Figure 8.** Principal component analysis (PCA) of gene expression. DESeq2 normalized counts were log-plus-one transformed and scaled to zero mean and unit variance prior to PCA decomposition. (A) PCA of all samples, labeled by sample type and study. (B) PCA of selected samples. Note that the samples appear grouped by study and not by cell type; see in particular Yamashiro *et al.* hiPSCs (brown squares), Tang *et al.* hESCs (green crosses), and hiPSCs from this study (blue circles).

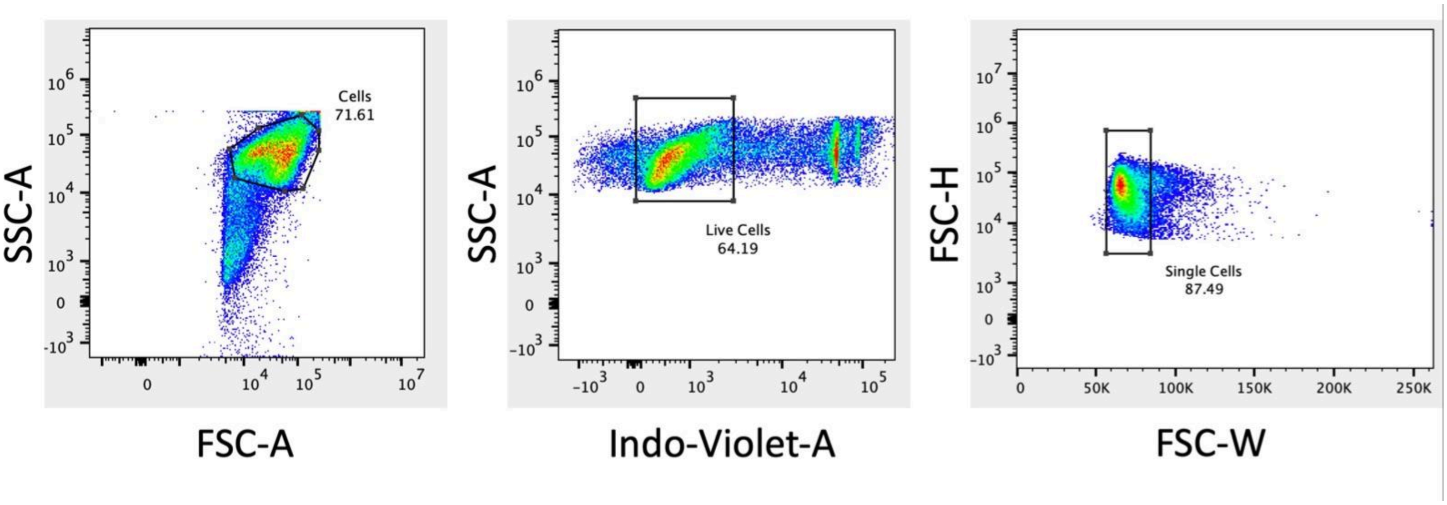


**Supplementary Figure 9:** Representative flow cytometry analysis, **Related to Figure 2, S4, and S5** A) Gating strategy for identifying cell events from debris using SSC-A versus FSC-A dispersion (left panel), live cells from dead cells using SSC-A by Indo-Violet-A dispersion in DAPI stained samples (center panel), and live single cells using FSC-H by FSC-W dispersion (right panel). A minimum of 10,000 live singlets were used for analysis. Analysis was performed in Flowjo software.
